# A strategy to address dissociation-induced compositional and transcriptional bias for single-cell analysis of the human mammary gland

**DOI:** 10.1101/2021.02.11.430721

**Authors:** Lisa K. Engelbrecht, Alecia-Jane Twigger, Hilary M. Ganz, Christian J. Gabka, Andreas R. Bausch, Heiko Lickert, Michael Sterr, Ines Kunze, Walid T. Khaled, Christina H. Scheel

## Abstract

Single-cell transcriptomics provide insights into cellular heterogeneity and lineage dynamics that are key to better understanding normal mammary gland function as well as breast cancer initiation and progression. In contrast to murine tissue, human mammary glands require laborious dissociation protocols to isolate single cells. This leads to unavoidable procedure-induced compositional and transcriptional bias. Here, we present a new strategy on how to identify and minimize systematic error by combining different tissue dissociation strategies and then directly comparing composition and transcriptome of isolated cells using single-cell RNA sequencing and flow cytometry. Depending on the tissue isolation strategy, we found dramatic differences in abundance and heterogeneity of certain stromal cells types. Moreover, we identified lineage-specific dissociation-induced gene expression changes that, if left unchecked, could lead to misinterpretation of cellular heterogeneity and, since the basal epithelial population is particularly affected by this, wrongful assignment of putative stem cell populations.

## Introduction

The mammary gland is a complex and heterogeneous organ consisting of an epithelial branched ductal network embedded in an extracellular matrix (ECM) containing a variety of different stromal cell types including fibroblasts, adipocytes, immune and endothelial cells. In both the human and murine mammary gland, the epithelial network consists of two main cellular compartments, an outer layer of basal/myoepithelial cells surrounding an inner layer of luminal cells. Control of cell fate and tissue function in the mammary gland takes place in the context of constant remodelling under the influence of steroid hormones and remains a topic of intense research^1,2^. Although breast cancer arises from the epithelial compartment, it is recognized that progression is driven through complex bi-directional interactions between tumour and stromal cells^3,4^. Therefore, unravelling mammary cellular heterogeneity within the epithelial and the stromal compartment together with a comprehensive understanding of epithelial cell fate conversions is essential not only to better understand mammary gland function, but is also of fundamental importance for many aspects of breast cancer initiation and progression.

Cellular heterogeneity within the two main epithelial populations has already been described. Specifically, in the human mammary gland two luminal subpopulations have been identified based on the markers EpCAM and CD49f, namely luminal mature (LM, EpCAM^+^/CD49f^−^) and luminal progenitor (LP, EpCAM^+^/CD49f^+^) cells^5^. Recent advances in single cell technologies, such as single cell transcriptomics, have enabled a more detailed examination of mammary cell heterogeneity and, in the murine mammary gland, have enabled the reconstruction of developmental trajectories. A growing number of single-cell RNA sequencing (scRNA-seq) studies have analysed human^6,7^ and mouse mammary epithelial cells (MECs)^8–12^ and indicated the presence of three main epithelial cell types, namely basal/myoepithelial cells and two luminal subsets that closely corresponded to the EpCAM^+^/CD49f^+^ and EpCAM^+^/CD49f^−^ population designated as LP and LM cells in the human mammary gland. Importantly, additional heterogeneity within these main epithelial populations was described, albeit not consistently between these studies. For example, one study identified two subclusters of basal cells within the human mammary gland that differed in the expression levels of myoepithelial marker genes, such *ACTA2* and similarly, distinct luminal cell states^6^.

Importantly, in contrast to the murine mammary gland, human breast tissue samples at specific developmental stages, such as lactation or prepubertal, are largely inaccessible for obvious ethical reasons. Hence, functional verification of scRNA-seq findings is limited to transplantation experiments and *in vitro* studies. These limitations, together with the need for reliable cross-study comparison of single-cell transcriptomic data reinforce the importance of addressing bias introduced by variations in tissue processing strategies and the isolation of single cells. For human breast tissue, isolation of single cells typically involves long mechanical and enzymatic dissociation where a majority of well-established protocols rely on a 16- to 24-hour enzymatic dissociation with varying concentrations of collagenase and/or hyaluronidase (Table S1) followed by trypsin and/or dispase treatment. During these processes, exposure to shear stress and enzymatic action can lead to the disruption of cell surface markers, widespread transcriptional changes, induction of apoptosis and, in the worst case, complete loss of certain cell subtypes^13–15^. Indeed, it has recently been shown that enzymatic tissue dissociation at a physiological temperature of 37°C can induce the upregulation of stress response genes^14–16^. In addition, a dissociation-induced subpopulation has been reported for murine satellite cells^17^, underscoring that dissociation-induced gene expression artefacts may be present in other scRNA-seq data sets, possibly affecting their interpretation.

Here, we analysed the influence of tissue dissociation parameters on isolated human mammary cells via flow cytometry and scRNA-seq and thereby identified sources of dissociation-induced transcriptional and compositional bias. By comparing three enzymatic dissociation protocols related to commonly used protocols with variable agitation speed and duration, we found that a more vigorous agitation speed of 100 rpm compared to 10 rpm affected the composition of isolated cells, specifically leading to a reduction in stromal cells, in particular fibroblasts, and endothelial cells. In addition, our study revealed transcriptional changes induced during a long, 16-hour dissociation protocol compared to a shorter, 3-hour protocol, which were characterized by induction of an oxidative stress response and downregulation of lineage-specific marker genes. Thereby, downregulation of the key-myoepithelial gene *ACTA2*, yielded two basal/myoepithelial subpopulations distinguishable by expression levels of *ACTA2*. In conclusion, we present a systemic analysis of dissociation-induced compositional and transcriptional bias in human breast tissue samples. The direct comparison of dissociation procedures provides a strategy to capture the complex composition as well as the primary transcriptional state of human mammary cell populations and thereby helps to delineate cellular heterogeneity and to avoid misinterpretation caused by exogenously introduced bias.

## Results

### Enzymatic dissociation protocols for human reduction mammoplasties differ in yield, but not viability

Various aspects of human breast tissue dissociation are likely to influence composition and viability of isolated mammary cells. The majority of dissociation protocols involve an enzymatic digest using collagenase and hyaluronidase, but vary in duration and agitation speed (Table S1). Importantly, such parameters have previously been shown to impact the FACS profile of isolated mammary cells^13,18^. In order to examine in depth how variations in the enzymatic dissociation protocol impact yield, viability and composition of isolated cells, we collected human reduction mammoplasties from healthy women (Table S2), mechanically dissociated the tissue and subsequently subjected the samples to three different enzymatic dissociation protocols related to commonly used protocols and varying in duration (3 hours vs. 16 hours) and in agitation speed (10 rpm vs. 100rpm) (Fig.1A). As previously described, the enzymatic dissociation process was considered successful when the digestion solution was free of visible tissue pieces and contained cellular fragments of ductal or alveolar morphology with no attached extracellular matrix (ECM)^19^ (Fig.S1A,B). Thus, protocol A consisted of a short, 3-hour enzymatic digest with vigorous shaking at 100 rpm and a collagenase concentration of 800 U/mL. At a lower collagenase concentration of 300 U/mL, a complete tissue digest could not be achieved within 3 hours, indicated by remaining tissue clumps in the digestion solution (data not shown). Incomplete tissue digest was also observed when lowering the agitation speed during a 3-hour dissociation from 100 rpm to 10 rpm (data not shown). Together, these findings suggest that a higher agitation speed and enzyme concentration are necessary to complete tissue dissociation in 3 hours. On the other hand, we found that for a long 16-hour protocol (protocol B and C), effective dissociation of mammary tissue was achieved at both an agitation of 100 rpm and 10 rpm (Fig.S1A). For the longer enzymatic dissociation protocols, a final concentration of 300 U/mL collagenase was used, as a higher collagenase concentration of 800 U/mL led to loss of CD49f/Integrin alpha 6 (Fig.S1C), an important cell surface marker for flow cytometry of MECs^5^. Cellular fragments obtained after enzymatic dissociation showed great heterogeneity in size and morphology (Fig.1B). For each protocol, we observed small round and stick-like cell clusters, larger ductal structures that intermittently showed side branches, as well as medium sized fragments with ductal and/or alveolar morphology (Fig.1B). As it is conceivable that duration and agitation speed could determine fragment size, we analysed this by measuring the area using brightfield images. However, we determined comparable size distributions of fragments for all three protocols (Fig.1C). Independently of size and morphology, immunofluorescence and confocal microscopy identified those fragments as largely epithelial with CK18^+^ (cytokeratin 18) luminal cells surrounded by αSMA^+^ (alpha smooth muscle actin) basal/myoepithelial cells (Fig.1D). While larger fragments showed a uniform outer layer of basal/myoepithelial cells, smaller epithelial cell clusters often lacked complete basal/myoepithelial coverage (Fig.1D). To analyse cell viability directly after enzymatic dissociation, a single-cell suspension was generated via trypsinization and subjected to flow cytometry-based cell viability analysis. Remarkably, all enzymatic dissociation protocols yielded highly viable cells (>95%) (Fig.1E, S1D). Experimental designs often require cryopreservation of isolated mammary cells. Accordingly, cells can either be frozen directly after the enzymatic dissociation as a mixture of single cells and cellular fragments or subjected to trypsinization first and then frozen as a single cell suspension (Fig.S1E). The latter resulted in a marked decrease in cell viability of up to 50% after thawing. By contrast, consistently higher cell viabilities (>90%) were obtained if trypsinization was performed after thawing (Fig.1F, S1E). This is likely due to the loss of cell-cell contacts in single-cell suspensions which are maintained in mammary fragments and have been found to help suppress apoptosis^20^.

**Figure 1:**
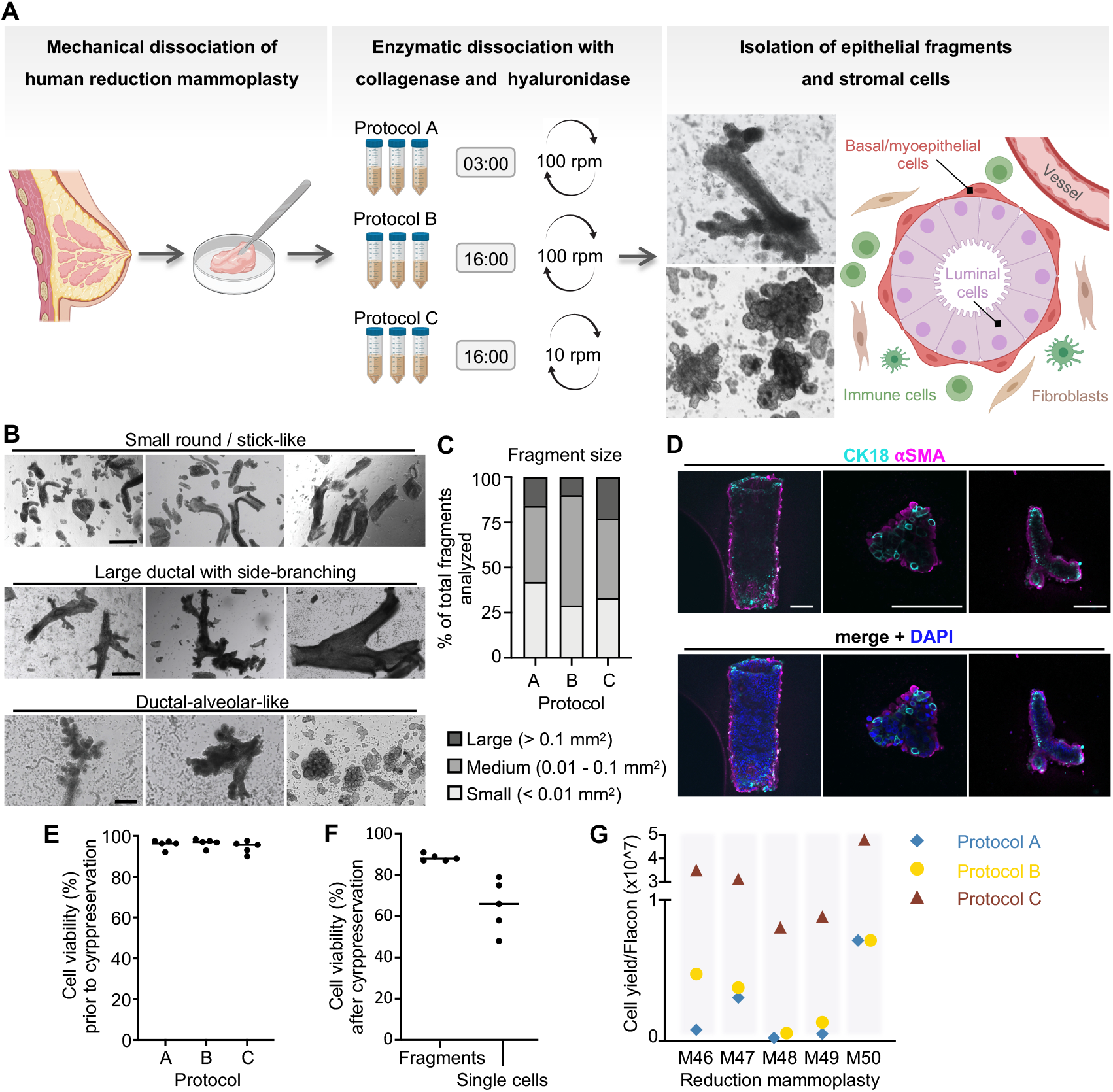
Differences in enzymatic dissociation protocols of healthy human breast tissue samples lead to discrepancies in cell yield, but not viability. **(A)** Following mechanical dissociation, human breast tissue is distributed over 50 mL falcons and subjected to three enzymatic dissociation protocols (protocol A, B, C) varying in agitation speed (100 and 10 rpm) and duration (3 and 16 hours) to yield epithelial fragments and stromal cells. Created with BioRender.com. **(B)** Brightfield images of cellular fragments after enzymatic dissociation. Reduction mammoplasties: M38, M47, M48, M50. Scale bar = 500 μm (upper two rows), 200 μm (lower row). **(C)** Size of cellular fragments after enzymatic dissociation. Total number of fragments analysed = 1228 (425 protocol A, 400 protocol B, 403 protocol. Reduction mammoplasties: M48, M49, M50. **(D)** Immunostaining of epithelial fragments embedded in collagen showed polarized expression of basal (aSMA) and luminal (CK18) lineage markers. Reduction mammoplasties: M48, M49. Scale bar = 50 μm. **(E,F)** Cell viability of isolated cells was determined via flow cytometry using live-dead marker 7-AAD **(E)** prior to cryopreservation. Reduction mammoplasties: M46, M47, M48, M49, M50 and **(F)** after cryopreservation as single cells and cellular fragments. Reduction mammoplasties: M30, M38, M43, M44, M45. **(G)** Cell-yield after enzymatic dissociation determined per 50 mL flacon of dissociation suspension. Reduction mammoplasties: M46, M47, M48, M49, M50. See also Figure S1.

### Epithelial composition is comparable across different isolation protocols whereas stromal populations are most efficiently derived at low agitation speed

In contrast to viability, we observed large differences in the number of cells yielded between the three enzymatic dissociation protocols. Specifically, in line with previous reports^13^, long and slow tissue dissociation (protocol C) yielded approximately 10 times more cells compared to both short (protocol A) as well as long dissociation with high agitation speed (protocol B, Fig.1G). These findings led us to question whether certain mammary cell subpopulations are preferentially isolated depending on agitation speed and duration of tissue dissociation. To delineate subpopulations of cells yielded from different enzymatic dissociation protocols, we performed flow cytometry analysis utilizing commonly used markers for human mammary cell identification^5^. After excluding doublets and dead cells (Fig.S2A), CD31 and CD45 were used to identify endothelial and hematopoietic cells (Stroma1 = CD31^+^/CD45^+^). Using EpCAM and CD49f on CD31^−^/CD45^−^ cells as previously described, yielded another stromal component, mainly consisting of fibroblasts (Stroma2 = EpCAM^−^/CD49f^−^) as well as three major epithelial populations: EpCAM^high^ luminal cells, consisting of a luminal progenitor (EpCAM^high^/CD49f^+^) and a luminal mature (EpCAM^high^/CD49f^low^) subpopulation, as well as the EpCAM^−/low^/CD49f^+^ population, containing basal/myoepithelial cells^5^ (Fig.2A). Previous work from our laboratory has shown that use of membrane-bound metalloprotease CD10 as an additional marker is essential to purify basal/myoepithelial cells from the EpCAM^−/low^/CD49f^+^ population^21^. Importantly, CD10 has an identical expression pattern as the key lineage marker alpha-smooth muscle actin (*α*-SMA) in basal/myoepithelial cells^22–26^. Cells that are EpCAM^−/low^/CD49f^+^/CD10^−^, were denoted as Stroma3 (Fig. 2A), and were previously shown to express lymph endothelial as well as immune cell markers^21^. Consistently across all 5 donors, we found that an agitation speed of 100 rpm (protocol A and B) yielded higher proportions of epithelial (basal and luminal) populations (Fig. 2B, S2B; Suppl. file 1). On average, 60% of isolated cells from protocol A and 85% from protocol B were epithelial, while an agitation speed of 10 rpm (protocol C) lead to increased proportions of stromal fractions (>80%), which was particularly striking for the Stroma2 and for the EpCAM^−/low^/CD49f^+^/CD10^−^ Stroma3 population. Whereas the EpCAM^−/low^/CD49f^+^ population in protocol A and B consisted mainly of CD10^+^ basal/myoepithelial cells (>85%) and only a small fraction of CD10^−^ cells, this ratio was reversed for protocol C (Fig. 2B, Fig.S2B, Suppl. file 1). Upon calculating absolute cell numbers per falcon of digested breast tissue for each subpopulation, we found that whilst the number of epithelial cells isolated were comparable between different dissociation protocols (Fig.2C), the number of overall stromal cells was much higher for protocol C (Fig.2D). Thus, considering the higher number of total cells isolated by protocol C, we concluded that although all protocols yield similar numbers of epithelial cells, a low agitation speed additionally yields a high number of stromal cells, particularly fibroblasts which reside in the EpCAM^−^/CD49f^−^ Stroma2 population and cells of the EpCAM^−/low^/CD49f^+^/CD10^−^ Stroma3 population, previously identified as a mixture of lymph endothelial as well as immune cells ^21^. Conversely, at a higher agitation speed of 100 rpm (protocol A and B), the total number of stromal cells is drastically decreased, skewing the composition towards the epithelial fractions, especially after a 16-hour dissociation at 100 rpm (protocol B).

**Figure 2:**
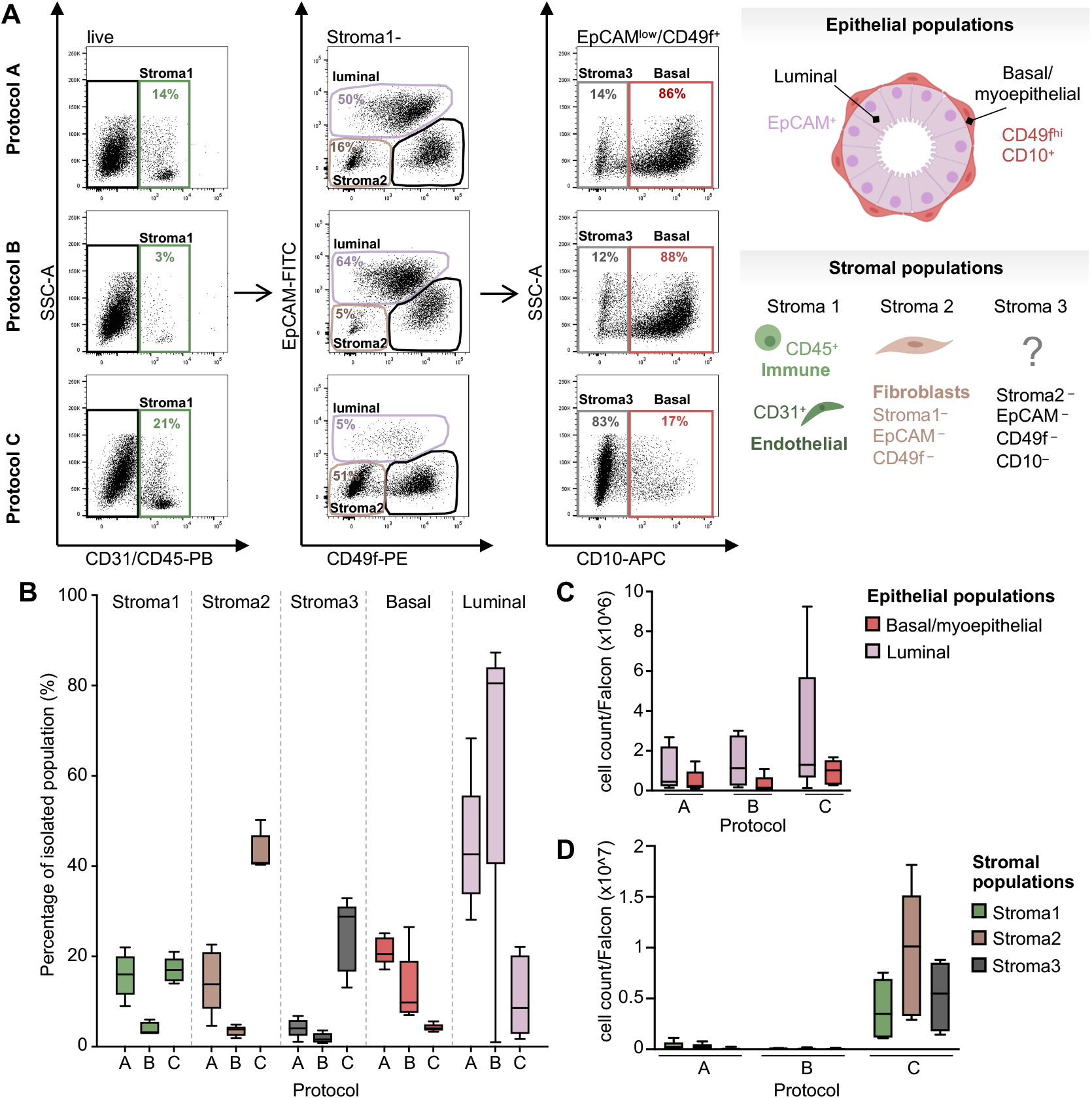
Epithelial cells are uniformly isolated across different isolation protocols, whilst stromal cells are preferentially isolated at low agitation speeds. **(A)** Flow cytometry analysis of mammary cells after enzymatic dissociation protocol A, B and C after exclusion of doublets and dead cells (live = 7-AAD^−^): CD31 and CD45 were used to depict the stroma1 population (CD31^+^/CD45^+^). Among the remaining CD31^−^/CD45^−^ cells, EpCAM and CD49f were used to delineate the following populations: stroma2 (CD49f^−^/EpCAM^−^), luminal (EpCAM^+^) and CD49f^+^/EpCAM^−/low^. The latter was further subdivided into stroma3 (CD10^−^/CD49f^+^/EpCAM^−/low^) and basal/myoepithelial (CD10^+^/CD49f^+^/EpCAM^−/low^). Reduction mammoplasty shown: M46. Created with BioRender.com. **(B)** Proportions of stromal and epithelial populations isolated with enzymatic dissociation protocols A, B and C based on flow cytometry analysis as shown in Fig.2SA and Fig.2A. Reduction mammoplasties: M46, M47, M48, M49, M50. **(C, D)** Total cell count of epithelial **(C)** and stromal **(D)** cells isolated with enzymatic dissociation protocols A, B, and C per 50 mL falcon of dissociated breast tissue. Reduction mammoplasties: M46, M47, M48, M49, M50. See also Figure S2.

### Single-cell RNA sequencing reveals dissociation-induced cellular differences

Analysis of cell surface markers by flow cytometry analysis, revealed that enzymatic dissociation protocols A, B, and C yielded different proportions of epithelial and stromal fractions. However, flow cytometry is limited by low multiplexing ability and not suitable for delineating greater heterogeneity within the main mammary cell populations, a problem overcome by single-cell transcriptomic approaches. Thus, in order to further specify the compositional differences in isolated cells and to examine whether different enzymatic dissociation protocols influence the cellular transcriptome, we performed scRNA-seq using the 10X Chromium platform of one reduction mammoplasty sample digested using protocols A, B, and C. Single-cell suspensions were generated from thawed fragments of each protocol and sequenced in one run on separate lanes after dead cell exclusion. After filtering out poor quality cells, we performed downstream analysis on 3,586 cells from protocol A, 2,809 cells for protocol B and 4,796 cells for protocol C with an average of 7,427 unique molecular identifiers and 2,445 genes detected per cell (Fig. S3A).

By plotting the Uniform Manifold Approximation and Projection (UMAP) dimensionality reduction, we identified 12 cell clusters (C1-C12, Fig. 3A). Interestingly, some of the identified clusters consisted of cells derived from both 16-hour dissociation protocols (C2, C4, C5, protocol B and C), while other clusters originated only from the 3-hour dissociation (C1, C3, protocol A, Fig. 3B). The remaining clusters contained cells from all three dissociation protocols (C6-C12, Fig. 3B, C). Thus, we observed that cells derived from the 16-hour dissociation (protocol B and C) formed largely overlapping clusters, while cells derived from the 3-hour protocol (protocol A) showed shifts in UMAP coordinates (Fig. 3B; S3B, C), suggesting transcriptional differences originating from the duration of the dissociation process. Next, we aimed to identify mammary cell types and therefore interrogated the expression of previously identified marker genes within the clusters. This analysis indicated the presence of three epithelial populations for each dissociation protocol: basal/myoepithelial (BA) cells and two luminal populations that closely corresponded to the EpCAM^high^/CD49f^+^ and EpCAM^high^/CD49f^low^ population designated here as luminal hormone-receptor negative progenitors (LHR^−^) and luminal hormone-receptor positive mature cells (LHR^+^). Two separate clusters (C1 from protocol A; C2 from protocol B/C) expressed known basal/myoepithelial lineage markers such as *ACTA2*, *OXTR* and *TP63* whereas four clusters (C3-C6) displayed high levels of the luminal marker KRT18. Specifically, clusters C6 showed expression of hormone receptors (*PGR/ESR1*) and markers such as amphiregulin (*AREG*) indicative of LHR^+^ cells, and three clusters (C3 from protocol A; C4 and C5 from protocol B/C) expressed the gene for aldehyde dehydrogenase 1A3 (*ALDH1A3*) which is a known progenitor marker ^27^ and indicative of LHR^−^ cells (Fig. 3A,C; S3D). Besides these epithelial populations, we identified four stromal cell populations, namely fibroblasts (FB, cluster C7), vascular accessory cells (VA, cluster C8), endothelial (ED, cluster C9) and immune cells (IM, clusters C10-C12) contributed to by all digestion protocols (Fig. 3A-C; S3D). Fibroblasts were classified based on expression of collagens, including *COL1A1*, and extracellular matrix protein genes such as *DCN* and *LUM* (Fig. 3C; S3D). Comparable to the basal/myoepithelial cells, vascular accessory cells expressed myoepithelial markers, like *ACTA2* and *TAGLN*. Additionally, they showed high levels of *GJA4*, a component of gap junctions (Fig.3C; S3D), and *ESAM*, an adhesion molecule that vascular accessory cells share with endothelial cells (Fig. 3C). Known endothelial lineage marker *PECAM1*, encoding for CD31, and *SELE*, a cell surface glycoprotein, were used to specifically identify endothelial cells (Fig. 3C; S3D). The immune cell population was characterized by expression of *PTPCR* which encodes for the immune cell marker CD45 and consisted of three different clusters (C10-C12), reflecting immune cell heterogeneity (Fig. 3C; S3D).

**Figure 3:**
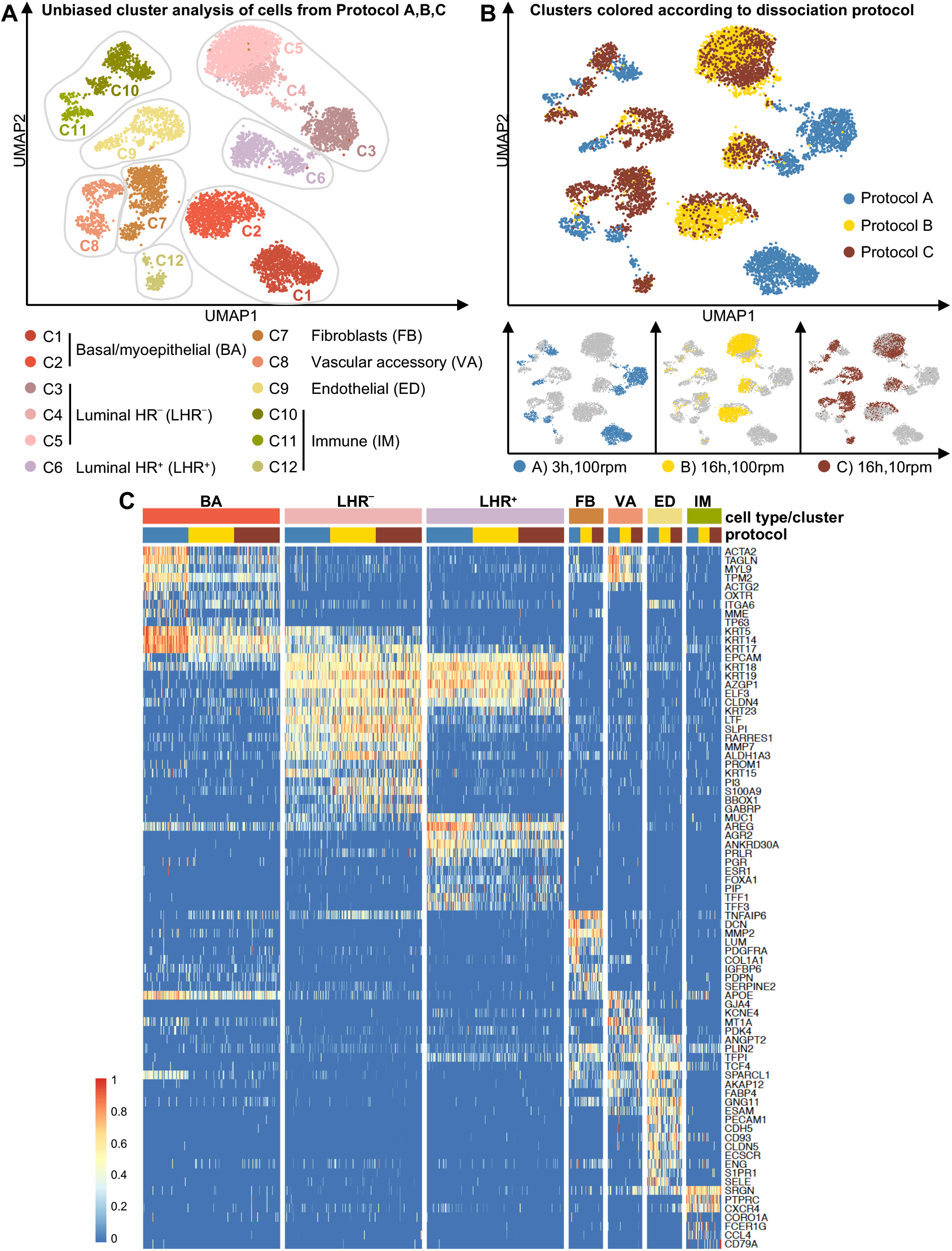
Single-cell RNA sequencing analysis of cells isolated with different enzymatic dissociation protocols. **(A)** 8,100 randomly sampled cells from dissociation protocol A, B and C of reduction mammoplasty M46 (2,700 cells per dissociation protocol) were visualized as 12 clusters in a UMAP plot and coloured by cluster. **(B)** Same as in (A) but cells were coloured by dissociation protocol A, B and C. **(C)** Heatmap showing key marker genes that were used to identify putative identities of mammary cell clusters. Upper bars represent cell type/cluster and dissociation protocol origin of cells. From each dissociation protocol, 100 randomly selected cells per sample of the epithelial clusters and 25 cells per sample of the stromal clusters are displayed for better visualization. Colour scale represents log-transformed and normalized counts scaled to a maximum of 1 per row. See also Figure S3.

Importantly, the compositional differences between the dissociation protocols, as already detected by flow cytometry analysis (Fig. 2), were also largely reflected by the scRNA-seq data. Comparable to the flow cytometry analysis an agitation speed of 100 rpm (protocol A and B) led to a higher representation of epithelial compared to stromal fractions (> 70%). However, we observed that the proportion of stromal compared to epithelial fractions isolated at a low agitation speed (protocol C) was approximately 90% according to flow cytometry data compared to 60% according to scRNA-seq data (Fig.S3E). One reason behind this discrepancy could be the downregulation or loss of cell surface markers caused by a long enzymatic dissociation periods, as we observe for a number of other lineage markers by scRNA-seq and describe in detail below.

Taken together, based on commonly used lineage markers, we identified mammary stromal and epithelial cell types comparable to previous studies^6,7^. However, by directly comparing different tissue dissociation protocols, we detected shifts between UMAP clusters originating from the 16-hour protocols compared to the 3-hour protocol (Fig.3B). This observation was most pronounced for the epithelial compared to stromal fractions, suggesting widespread transcriptional changes due to the duration of enzymatic dissociation. Based on these observations, we first sought to further delineate the rich stromal heterogeneity yielded by protocol C and then detail the transcriptomic changes caused by duration of enzymatic dissociation.

### Representation of stromal heterogeneity depends on agitation speed during enzymatic tissue digestion

Flow cytometry and scRNA-seq data both showed an enrichment for stromal fractions at a lower agitation speed of 10 rpm (protocol C). As a next step, we wished to further examine cellular heterogeneity within the stromal compartment. We were also interested to determine which stromal populations were specifically enriched or reduced as a consequence of the dissociation protocol applied. For this purpose, we subdivided the stromal clusters (C7-C12; Fig.3A) into 12 subclusters (SC1-12, Fig. 4A, S4A). In general, subdivision of the three immune cell clusters (C10-C12; Fig.3A) revealed five subclusters (SC8-12, Fig.4A) containing a mix of cells derived from different protocols (Fig. 4B). These data suggested that each immune cell type is represented for each dissociation protocol, although in different numbers, and no large transcriptomic changes occurred as a result of the long enzymatic digest. By contrast, the clusters containing fibroblasts, vascular accessory cells and endothelial cells (C7-C9; Fig.3A, respectively) fell into 7 subclusters that were distinguishable by differences in both duration and agitation speed of enzymatic dissociation (SC1-7, Fig.4A).

**Figure 4:**
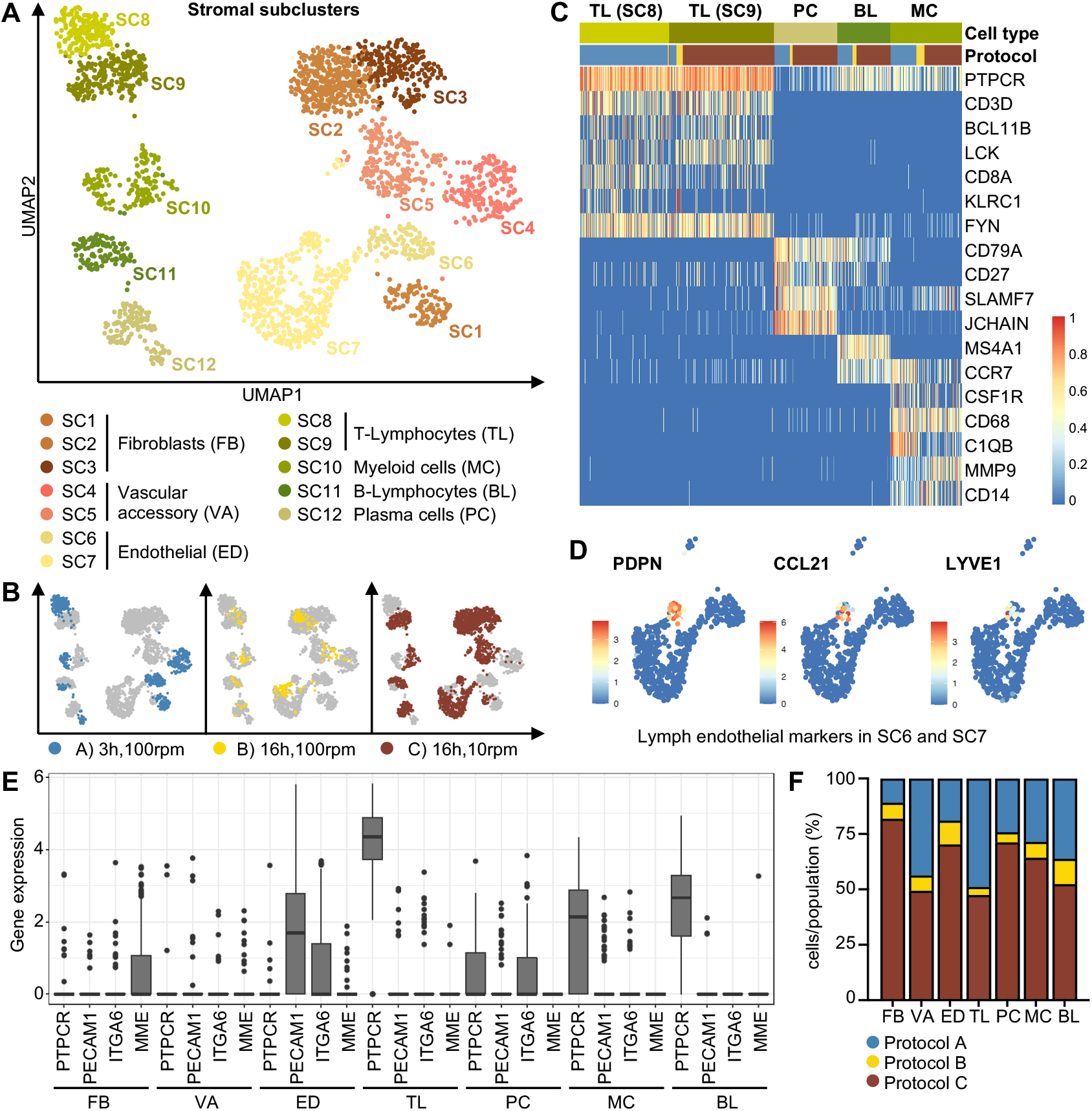
Agitation speed determines stromal heterogeneity. **(A)** Stromal cell clusters identified in Fig. 3 were re-clustered, yielding 12 subclusters (SC1-SC12), which were visualized in a UMAP plot. **(B)** Same as (A) but coloured by dissociation protocol A, B and C. **(C)** Heatmap showing key marker genes used for identification of immune cell clusters. Upper bars represent cell type/cluster and dissociation protocol origin of cells. All cells of the immune cell subclusters (SC8-SC12) derived from the 3-hour (protocol A) and the 16-hour dissociation (protocol B and C) are displayed. Colour scale represents log-transformed and normalized counts scaled to a maximum of 1 per row. **(D)** UMAP plots showing the endothelial subclusters SC6 and SC7 coloured by normalized log-transformed expression of lymphatic marker genes *PDPN*, *CCL21* and *LYVE1*. **(E)** Box plot for expression of *PTPCR*, *PECAM1*, *ITGA6*, and *MME* by cell type showing expression of *ITGA6* in endothelial (ED) and plasma cells (PC). **(F)** Proportions of cells derived from dissociation protocol A, B, and C identified as stromal cell type. See also Figure S4.

Marker analysis of the 5 immune cell subclusters revealed expression patterns indicative of T-lymphocytes characterized by high expression of *CD3D* (TL, subcluster SC8 and SC9), *CD79A* positive B-lymphocytes (BL, subcluster SC11) and myeloid cells (MC, subcluster SC10) expressing macrophage and monocyte markers *C1QB* and *CD68* (Fig. 4A, C; S4B). Plasma cells were identified based on expressing the IgA and IgM protein component encoded by *JCHAIN* and were negative for *MS4A1*, which encodes for *CD20* and is typically downregulated in plasma cells after differentiation from B-lymphocytes (PC, subcluster SC12, Fig. 4A,C; S4B) ^28,29^. Taken together, these results indicated that each immune cell subcluster was generally attributable to a certain cellular subtype: T-lymphocytes, plasma cells, B-lymphocytes and myeloid cells.

With respect to the remaining clusters, digestion duration (either 3 or 16 hours) and agitation speed (either 100 or 10 rpm) appeared to generate subclusters arising in fibroblasts (SC1-SC3), vascular accessory cells (SC4/5), and endothelial cells (SC6/7) (Fig.4A). For those subclusters, we noted that protocol C (16h, 10rpm) yielded five subclusters (SC1-3, SC5, SC7), while protocol A (3h, 100rpm) and B (16h, 100rpm) only yielded three subclusters: SC1, SC6, SC4 for protocol A and SC2, SC5, SC7 for protocol B (Fig.4A, B). This increased stromal heterogeneity for protocol C was particularly striking for the population of fibroblasts. More than 80% of fibroblasts were isolated with protocol C giving rise to three fibroblasts clusters (SC1-3), while protocol A and B yielded less fibroblasts that clustered in only one subcluster each (SC1 for protocol A, SC2 for protocol B) (Fig.4A, B). Further distinctions could be found in the marker expression of endothelial cells apparently caused by the agitation speed applied. Specifically, within the endothelial cell subcluster arising mostly from protocol C (SC7), expression of known lymphatic markers like *PDPN*, *CCL21* and *LYVE1* could separate the endothelial cells further into either vascular or lymphatic endothelial cells (Fig. 4D). As described earlier, lymphatic endothelial cells have previously been detected in the EpCAM^−/low^/CD49f^+^/CD10^−^ Stroma3 population identified via flow cytometry, a population of cells preferentially isolated at a low agitation speed (protocol C). When taking a closer look at the marker expression of endothelial cells from both subclusters SC6 and SC7, including the lymphatic endothelial cells, we detected expression of *ITGA6* which encodes for CD49f (Fig. 4E) and in line with previous reports, *LYVE1*^+^ lymphatic cells expressed *PECAM1* (encoding CD31), but only low levels ^30^ (Fig.S4C). Another cell population previously identified within the EpCAM^−/low^/CD49f^+^/CD10^−^ Stroma3 population were plasma cells. Interestingly, this cell type also expressed *ITGA6*, as has been reported before ^31^ (Fig.4E). Upon closer inspection, our data also revealed that plasma cells expressed rather low levels of *PTPCR* (encoding CD45), which is in line with the described progressive decline of CD45 during plasma cell differentiation ^32^.

In summary, our findings strongly indicated that the EpCAM^−/low^/CD49f^+^/CD10^−^ Stroma3 population identified via flow cytometry contains vascular and lymphatic endothelial cells as well as plasma cells, despite prior exclusion of such cells by CD31 and CD45. Thus, these data reinforce the importance of CD10 as a cell-surface marker to identify basal/myoepithelial cells by flow cytometry. Importantly, more than 70 % of the Stroma3 population were isolated at a low agitation speed of 10 rpm (protocol C). Together with the observation of increased fibroblast heterogeneity at a lower agitation speed, these results suggest protocol C as particularly suitable if isolation of large numbers of stromal cells and delineation of stromal heterogeneity is desired, particular with respect to fibroblasts and endothelial cells. Other stromal populations such as vascular accessory and immune cells were well-represented even when a higher agitation speed was applied for 3 hours (Protocol A). However, longer periods of high agitation (Protocol B) led to a reduction of all stromal cell types, thus strongly favouring low agitation in combination with long enzymatic digestion if an effective isolation of stromal fractions is desired (Fig. 4F).

### Longer enzymatic tissue dissociation leads to an oxidative stress response in mammary cells

As described above, differences in enzymatic dissociation duration led to discrepancies in the transcriptomic profile of epithelial and stromal subtypes indicated by shifts in representative clusters or even the generation of new clusters (Fig.3A, B). To investigate how the transcriptional state of isolated cells changed in response to increased dissociation duration, we generated an average gene expression score for cells isolated using the 16-hour (Protocol B and C) compared to the 3-hour (Protocol A) dissociation protocol, where cell subpopulation discrepancies were accounted for by firstly averaging across cell type, then normalizing within each dissociation duration group.

Differentially expressed genes were identified as having a fold change in expression greater than 1.5 between the 16-hour and 3-hour dissociation. Overall, we found a total of 192 genes differentially expressed. Specifically, 60 genes were found to be upregulated and 132 genes were downregulated after 16-hour compared to the 3-hour dissociation process (Fig. 5A; Suppl. file 2). Gene ontology (GO) term analysis of the 60 upregulated genes revealed an enrichment for enzymes with oxidoreductase activity indicative of a cellular response to oxidative stress (Fig.5B, S5A; Supplementary file 10). Included in these pathways were genes encoding detoxifying enzymes known to protect from oxidative stress such as microsomal glutathione S-transferase (*MGST1*)^33^, NAD(P)H quinone dehydrogenase 1 (*NQO1*) ^34^ and heme oxygenase 1 (*HMOX1*) ^35^. Similarly, *GCLM* was upregulated in cells isolated with the 16-hour dissociation protocols, which encodes for a subunit of the glutamate cysteine synthase, crucial for the synthesis of one of the most important cellular antioxidants, glutathione (GSH) ^36–38^. *TKT* and *TALDO* were also found to be upregulated and encode for rate limiting enzymes of the pentose phosphate pathway which produces NADPH, important in providing reducing potential to antioxidants and redox regulatory enzymes ^39–41^. Besides these findings, we detected the two subunits, *FTH1* and *FTL*, of iron storage protein ferritin, which helps to reduce accumulation of reactive oxygen species and thereby protects against iron-mediated lipid peroxidation ^42–45^.

**Figure 5:**
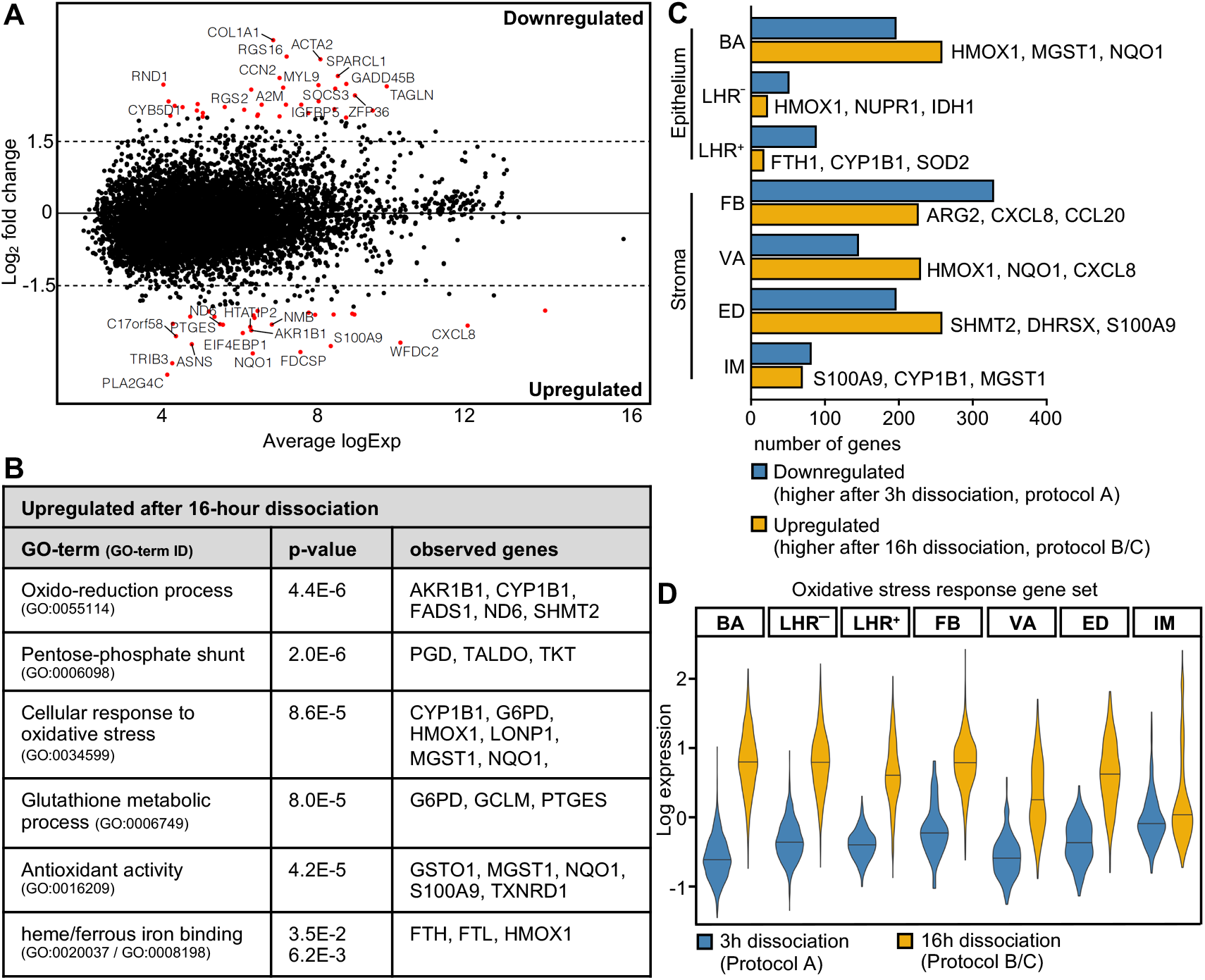
Long enzymatic dissociation induces an oxidative stress response. **(A)** MA plot showing fold changes for genes differentially expressed between cells derived from the 16-hour dissociation (protocol B and C) and cells derived from the 3-hour dissociation (protocol A). Genes with a log fold change of at least 1.5 were identified and notated in the graph coloured in red. 60 genes were higher expressed (upregulated) after a 16-hour dissociation (protocol B and C) and 132 genes after a 3-hour dissociation (downregulated, protocol A). **(B)** GO term analysis of 60 genes upregulated in cells after a 16-hour dissociation (protocol B and C) compared to a 3-hour dissociation (protocol A). Selected significantly enriched terms (P<0.01) are displayed and show an enrichment for oxidative stress response genes. **(C)** Bar graph showing number of differentially expressed genes per cell type (log fold change > 2.0). Selected genes displayed next to the bars were picked among the top upregulated genes. **(D)** Violin plot showing expression of oxidative stress response gene set: 14 genes selected among the upregulated genes after a 16-hour dissociation that play a role in cellular oxidant detoxification, GSH synthesis, iron metabolism and NADPH production and are associated with the corresponding GO terms. A list of genes together with the associated GO terms is displayed in table S3. Expression levels are displayed per cell type for the 3-hour (protocol A) and the 16-hour dissociation (protocol B and C). See also Figure S5.

To investigate whether tissue dissociation duration differentially impacted the transcription profile of mammary cell subpopulations, we performed individual differential gene expression analysis with a fold change cut off of 2.0 for each cell type isolated from the 16-hour compared to the 3-hour protocol. Thereby, we found that the number of upregulated genes after a 16-hour dissociation process differed between the cell types and was highest for basal/myoepithelial cells, fibroblasts, vascular accessory and endothelial cells, with an average of 244 genes upregulated with a log fold change of at least 2.0, compared to an average 37 genes upregulated in the two luminal LHR^+^ and LHR^−^ and the immune cell population (Fig. 5C; Supplementary files 3-9). Among genes most consistently upregulated across cell types we found again many enzymes with oxidoreductase activity and detoxifying genes (Fig. 5B, C). To further investigate whether certain cell populations are impacted more than others, we compiled a list of 14 genes involved in cellular oxidant detoxification, glutathione metabolism, iron metabolism and NADPH production found among the top upregulated genes after the 16-hour dissociation process and generated a stress response gene set (Table S3) that could be tested for enrichment within each mammary cell subtype. Analysis of the stress response gene set per cell type revealed that across all mammary subpopulations, except the immune cell population, cells isolated from the 16-hour protocols had higher levels of the stress response gene set (Fig. 5D), suggesting that protective mechanisms against dissociation-induced stress were activated systemically. Taking a closer look into cell-type specific expression of the stress response genes, we found that *MGST1* was upregulated specifically in the 16-hour digests for both epithelial populations and fibroblasts, whereas basal/myoepithelial and endothelial cells showed the strongest upregulation of *TXNRD1*, a member of the thioredoxin system and essential for redox homeostasis ^46,47^ (Fig. S5B,C). Thus, although, individual cell types showed slightly different levels of specific stress-response genes, we concluded that cells isolated from the 16 compared to 3-hour dissociation process undergo an overall oxidative stress response, presumably as a protection mechanism.

### Expression of lineage specific markers across different mammary cell subtypes is altered by the duration of enzymatic dissociation

Although genes upregulated in response to a 16-hour compared to 3-hour dissociation process were associated with a generalized cellular response to oxidative stress, GO term analysis of genes downregulated instead showed an enrichment for genes encoding for heat shock proteins, extracellular matrix constituents and binding proteins, as well as genes involved in immune receptor activity (Fig. 6A, S6A; Supplementary files 11-15).

**Figure 6:**
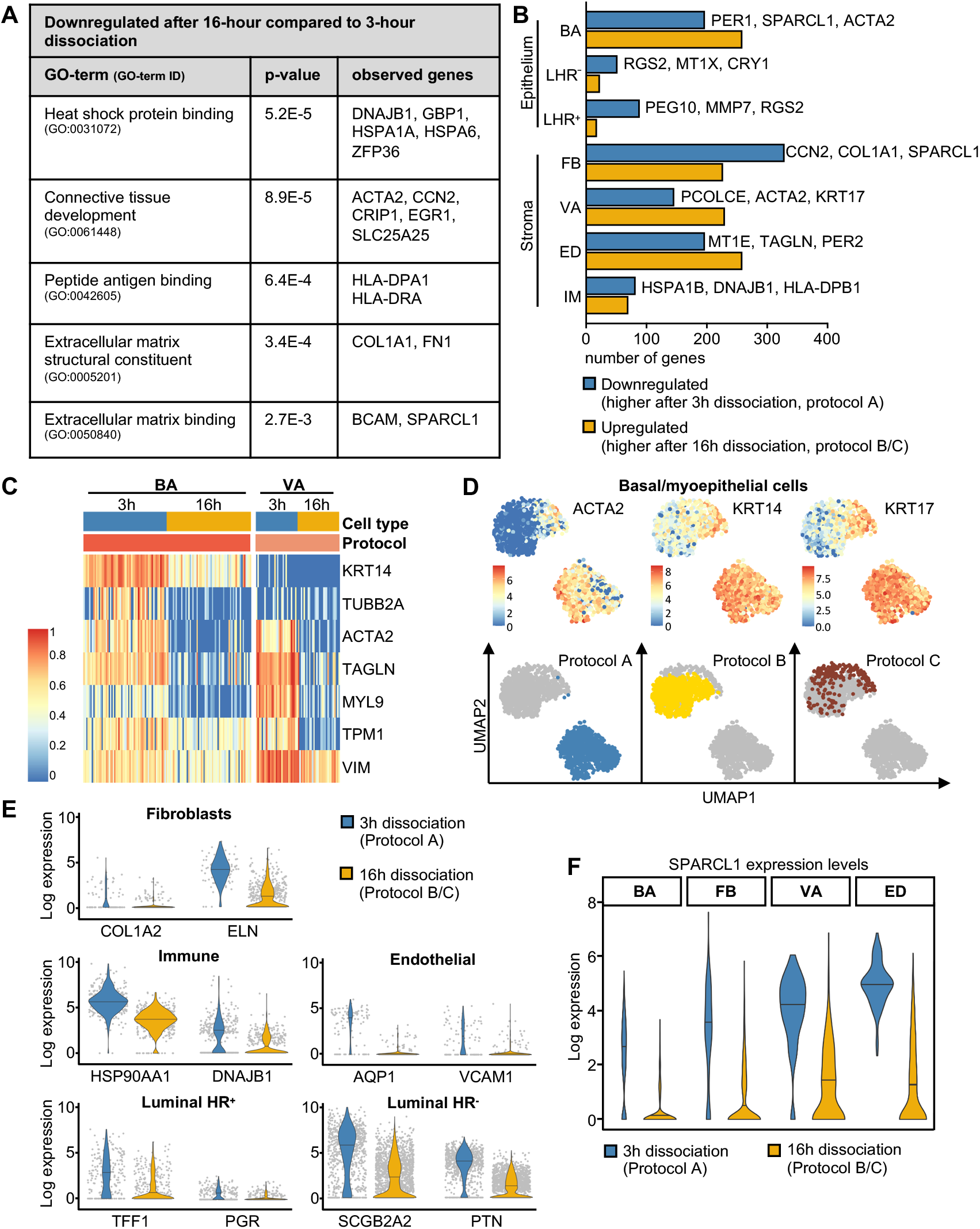
Downregulation of lineage specific markers after 16-hour protocols. **(A)** GO term analysis of 132 genes downregulated (log fold change > 1.5) in cells after a 16-hour dissociation (protocol B and C) compared to a 3-hour dissociation (protocol A). Selected significantly enriched terms (P < 0.01) are displayed. **(B)** Bar graph showing differentially expressed genes per cell type (log fold change > 2.0). Selected genes displayed next to the bars were picked among the top downregulated genes after a 16-hour dissociation. **(C)** Heatmap showing genes downregulated (log fold change > 2.0) in basal/myoepithelial and vascular accessory cells after a 16-hour (protocol B and C) compared to a 3-hour dissociation (protocol A) that were assigned to the GO term “structural constituents of cytoskeleton”. Upper bars represent cell type/cluster and dissociation protocol origin of cells, 3-hour (protocol A) and 16-hour dissociation (protocol B and C). 60 randomly selected basal/myoepithelial cells and 30 cells from vascular accessory for each dissociation duration group are displayed for better visualization. Colour scale represents log-transformed and normalized counts scaled to a maximum of 1 per row. **(D)** UMAP plots showing the basal/myoepithelial clusters C1 and C2 coloured by normalized log-transformed expression of marker genes *ACTA2*, *KRT14* and *KRT7* (upper row) and coloured by dissociation protocol origin (lower row). **(E)** Violin plots showing expression of selected genes found to be downregulated after a 16-hour (protocol B and C) compared to a 3-hour dissociation (protocol A) in fibroblasts (FB), immune cells (IM), endothelial cells (ED), and the two luminal populations, LHR^+^ and LHR^−^. Cells are displayed as grey dots. **(F)** Violin plot showing expression level of the gene *SPARCL1* in basal/myoepithelial (BA), fibroblasts (FB), vascular accessory (VA), and endothelial cells (ED) revealed downregulation after 16-hour (protocol B and C) compared to 3-hour dissociation (protocol A). See also Figure S6.

Comparable to the number of upregulated genes, the number of downregulated genes after a 16-hour dissociation varied among the different cell types with an average of 217 genes downregulated for basal/myoepithelial cells, fibroblasts, vascular accessory and endothelial cells and 74 genes for the two luminal (LHR^+^ and LHR^−^) and the immune cell population (Fig.6B; Supplementary files 3-9). A closer analysis of differentially downregulated genes revealed a number of cell-type specific changes in gene expression. Specifically, cells clustered as basal/myoepithelial and vascular accessory were among the mammary cell types that showed the highest number of downregulated genes after 16 hours of dissociation (Fig. 6B). Among the genes downregulated most strongly, many encoded for structural constituents of the cytoskeleton including certain keratins, like *KRT15* and *KRT17* and, importantly, known myoepithelial and vascular accessory lineage markers such as *ACTA2*, *TAGLN* and *MYL9* (Fig.6C, Fig.S6B). *ACTA2* encodes for alpha smooth muscle actin (*α*-SMA), a marker consistently expressed by basal/myoepithelial cells both in ducts and lobular acini ^26^. Closer analysis of *ACTA2* expression between the different dissociation protocols revealed a homogenous expression level among basal/myoepithelial cells after a 3-hour dissociation, whereas 16 hours of dissociation resulted in a heterogeneous population containing two basal subclusters with either high or low *ACTA2* levels (Fig.6D, Fig.S6C). Moreover, similar gene expression profiles were observed for genes encoding basal lineage marker keratins (*KRT14* and *KRT17*) (Fig.6D). These findings clearly suggest that longer enzymatic dissociation alters gene expression in a way that creates cellular heterogeneity within cells that have previously clustered tightly together, which is of particular concern when using this information to imply cellular lineages and hierarchies of the mammary epithelium.

Besides structural constituents of the cytoskeleton, we observed a downregulation of genes encoding for constituents of the extracellular matrix and genes involved in cell adhesion. Particularly, cells clustered as fibroblasts downregulated genes encoding for collagen (*COL3A1* and *COL1A1*), fibronectin (*FN1*), and elastin (*ELN*) (Fig.6E). Collagen fibril assembly levels have been shown to be regulated by the matricellular protein hevin, encoded by *SPARCL1* ^48^, which incidentally we found to be downregulated in several cell types including fibroblasts, endothelial, vascular accessory and basal/myoepithelial cells (Fig.6F). Of note, *SPARCL1* has been reported to act as tumour suppressor downregulated in many cancers including breast cancer ^49,50^.

Within the stromal population of endothelial and immune cells, downregulation of gene expression as a consequence of longer enzymatic dissociation also altered interpretation of cellular properties or identities. For example, differences in gene expression within the endothelial cell cluster revealed downregulation of aquaporin 1 (encoded by *AQP1*), which plays an important role in micro-vessel permeability, and the cell adhesion molecule VCAM-1 (encoded by *VCAM1*) which is expressed on both large and small blood vessels and mediates adhesion of immune cells to the endothelium ^51,52^ (Fig.6E). On the other hand, differences in gene expression within the immune cell cluster revealed downregulation of several genes encoding for heat shock proteins such as *HSP90AA1* and *DNAJB1*, which are involved in antigen presentation and activation of many immune cell subtypes (Fig.6E) ^53,54^. The two luminal populations also displayed downregulation of lineage specific genes; however, the number of downregulated genes was lower compared to the basal/myoepithelial population (Fig. 6B, E). Cells clustered as luminal HR^+^ downregulated the oestrogen target gene *TFF1*, and *PGR*, which encodes for progesterone receptor, a marker commonly used to distinguish between LHR^+^ and LHR^−^ cells present in the mammary epithelium ^55^ (Fig. 6E). Conversely, mammaglobin A (encoded by *SCGB2A2*) and lipophilin B (encoded by *SCGB1D2*), which both belong to the secretoglobin family, are known to be co-overexpressed in breast cancer ^56^ and here we found both genes strongly downregulated in luminal HR^−^ cells after a 16-hour compared to 3-hour dissociation. Besides that, we found *PTN* to be downregulated, which encodes for the heparin-binding growth factor pleiotrophin (Fig. 6E). *PTN* expression has previously been shown to decrease in mice mammary glands during lobular-alveolar differentiation around mid-pregnancy ^8,57^.

In summary, we demonstrate that breast tissue dissociation duration heavily impacts the gene expression profile of mammary cell populations, such as basal/myoepithelial, vascular accessory, fibroblasts and endothelial cells and, to a lesser extent, immune cells and the two luminal populations. Importantly, thereby, the expression of certain lineage specific marker genes was reduced as well as genes reported to be involved in cellular differentiation or cancer progression. Particularly for the basal/myoepithelial population, the dissociation-induced downregulation of key-myoepithelial lineage marker *ACTA2* may affect biological interpretation of cellular heterogeneity within this population. On the other hand, we clearly show that longer enzymatic dissociation at a low agitation speed (protocol C) may be desirable in instances where the research focus lies on analysis of the stromal compartment, particularly fibroblasts and endothelial cells, as these are derived in much greater abundance.

## Discussion

Rapid technical advancement in single cell transcriptomics offers new possibilities for unravelling cellular heterogeneity and differentiation dynamics in the mammary gland. However, the experimental steps necessary prior to sequencing potentially introduce pre-selection bias altering downstream scRNA-seq data analysis and interpretation. Especially for the human mammary gland and other tissues similarly rich in dense ECM that is not easily broken down, the generation of a high-quality, high-viability single-cell suspension represents a crucial step. Here, we applied flow cytometry and scRNA-seq to systematically describe alterations introduced by experimental manipulation that might alter cellular composition and, as a consequence, interpretations of heterogeneity, lineage identity and gene expression of cells presumed to represent an “*in situ*” or “primary state”.

Thus, we discovered that the composition and the amount of mammary epithelial cell populations was comparable and included all three major populations (basal/myoepithelial, luminal HR^+^ and luminal HR^−^) when enzymatic dissociation was performed either at low agitation speed of 10 rpm for 16 hours (protocol C) or at more vigorous agitation of 100 rpm for 3 (protocol A) or 16 hours (protocol B). However, applying 100 rpm for 16 hours (protocol B) drastically reduced the proportions of all stromal cell populations as detected via flow cytometry and scRNA-seq. Importantly, shortening of the dissociation period to 3 hours (protocol A) resulted in better representation, but still yielded significantly fewer stromal cells (roughly 40% of isolated cells) compared to protocol C (roughly 80% of isolated cells). This is particularly important for sequencing approaches that rely on sequencing moderate numbers of cells instead of thousands. At a lower agitation speed of 10 rpm (protocol C) roughly 10 times more stromal cells were isolated and importantly, scRNA-seq analysis revealed stromal subpopulations such as the lymphatic endothelial cells and an additional fibroblast subcluster that were not detected at a higher agitation speed (protocol A and B). We suspect that at a higher agitation speed, many stromal cell types suffer from the resulting shear stress, which leads to cell bursting and consequently to a loss of those cells, especially when this shear stress acts on the cells for a longer period of time (protocol B). These findings implicate protocol B as an unfavourable method to isolate and study mammary cells due to the loss or damage of stromal cells. Importantly, the stromal compartment is well known to drive breast cancer development ^58^ and many stromal cell types serve as biomarkers for various subtypes of breast cancer and correlate with prognosis ^59,60^. Consequently, stromal cell populations are increasingly targeted therapeutically, highlighting the importance of capturing full stromal heterogeneity during tissue processing steps.

Interestingly, our data also showed that cells clustered as plasma cells and certain endothelial cells such as lymph endothelial cells, expressed CD49f, encoded by *ITGA6*, a marker also used in combination with absent or low EpCAM expression to denote basal/myoepithelial cells. This finding suggests that the EpCAM^−/low^/CD49f^+^/CD10^−^ Stroma3 population, previously discovered by flow cytometry ^21^, indeed contains both endothelial and immune cells, despite prior exclusion of such cells by CD31 and CD45, which are not expressed or expressed at very low levels by these particular subtypes. Together, these results strongly suggest that CD10, which is ubiquitously expressed by basal/myoepithelial cells ^26^ is a highly useful additional marker to delineate basal/myoepithelial cells and also underscores the usefulness of combining cell surface marker analysis by flow cytometry with scRNA-seq to interpret cellular heterogeneity. Especially when using flow cytometry for the enrichment of basal/myoepithelial cells prior to sequencing, the surface marker CD10 is essential.

Besides compositional differences, we found that the duration of enzymatic dissociation impacted the transcriptome of isolated epithelial and stromal cell populations in a way that leads to misinterpretation of cellular heterogeneity and, possibly, lineage identity. This was particularly the case for the basal/myoepithelial population where a long dissociation process led to the downregulation of cell-type specific marker alpha smooth muscle actin, encoded by *ACTA2*. This downregulation induced heterogeneity with the basal subset where two basal subclusters with varying expression levels of *ACTA2* could be observed. Importantly, immunohistochemistry analysis of normal human breast samples has shown that alpha smooth muscle actin is consistently expressed by basal cells both in ducts and lobular acini ^26^ underscoring that the observed heterogeneity is an artefact induced by long dissociation duration unlikely reflecting differences that correspond to cell states observed *in situ*. Indeed, downregulation of *ACTA2* expression along with other markers was also observed in cells clustered as vascular accessory cells. Together, these results suggest this observation to represent a general response of contractile cell types to loss of anchorage and cell-cell contacts during tissue dissociation. Importantly, changes in gene expression were not limited to contractile cells. Downregulation of lineage-specific genes was observed for every mammary cell population identified by scRNA-seq. In addition, we found the expression of several cancer-associated genes, such as *SPARCL1* or mammaglobin A (encoded by *SCGB2A2*) to be affected by a long enzymatic dissociation process. Thus, especially when comparing data sets to identify cancer-associated or developmentally regulated genes, care must be taken to avoid misinterpretation.

Regarding the number of up- and downregulated genes after a 16-hour compared to a 3-hour dissociation procedure, the basal/myoepithelial cell population was more affected than the luminal cells. Basal/myoepithelial cells display a higher abundance of cell-matrix adhesion related proteins compared to luminal cells, which is in accordance with their function in the adhesion of the mammary epithelium to the basement membrane ^61,62^. The ECM in turn provides structural support to the epithelial ductal network, which is specifically targeted for destruction during enzymatic dissociation. As a consequence, basal/myoepithelial cells experience a loss of cell-ECM adhesion and we hypothesize that, as a result of transcriptional feedback, downregulate structural constituents of their cytoskeleton, coupled via cell-matrix adhesion proteins to the ECM ^62^. Likewise, other cell populations such as the vascular accessory cells and the fibroblasts display similar cell-matrix adhesion and accordingly, we identified these cells among the types most affected by a long enzymatic dissociation. Luminal populations, in turn, showed fewer transcriptional changes, which is in line with their inner, more protected position in the mammary epithelium. Within the epithelial fragments isolated from fresh breast tissue, we observed maintenance of the bilayered architecture of the mammary gland, assuring cell-cell contacts for the luminal cells with each other and with the myoepithelium. However, also for luminal populations, we discovered cell-type specific responses to a long dissociation duration. While LHR^+^ cells downregulated genes, involved in hormone signalling, LHR^−^ cells were found to downregulate several secretoglobin encoding genes, some of them reported to be deregulated during cancer progression.

Finally, our data revealed that epithelial as well as stromal populations respond to a 16-hour dissociation process with the upregulation of oxidative stress response genes. Importantly, transcriptional changes induced by proteolytic breakdown of tissue at 37°C have been reported before, and one strategy to minimize those artefacts involves a bacterial serine protease, which has been shown to digest various tissues including human breast cancer samples at 4 to 6°C ^14–16^. At lower temperatures, the transcriptomic machinery is less active and therefore, transcriptional changes are minimized. However, this dissociation strategy needs to be tested in order to determine its suitability in recovering the full cellular heterogeneity of the human mammary gland.

Our data clearly suggest that the dissociation duration must be kept to a minimum in order to preserve the “in situ” cellular gene expression state. On the other hand, a short tissue dissociation might be at the expense of yielding certain stromal subpopulations. Thus, we suggest that capturing the full cellular heterogeneity of mammary breast tissue and at the same time identifying dissociation-induced transcriptomic changes can be achieved by combining both dissociation strategies, a long, 16-hour enzymatic dissociation with gentle agitation and a short, 3-hour digest with more vigorous shaking. Irrespective of the tissue dissociation strategy, it is crucial to exhaust the techniques available for the analysis of human tissue, which is limited to *ex vivo* studies and is not amenable to approaches such as lineage tracing. Our study highlights the benefit of combining flow cytometry and in-depth transcriptional analysis for analysing tissue heterogeneity, not only for the purpose of enrichment of a certain population prior to sequencing but to better understand tissue composition. At present, scRNA-seq techniques are not easily scalable to a large number of donors, thus, flow cytometry provides an invaluable tool to obtain first indications of the extent of inter-donor variation. At the same time, it is important to develop computational tools to correct for unwanted technical artefacts allowing cross study comparisons ultimately leading to a robust understanding of human mammary subpopulations and their developmental potential.

## Experimental procedures

### Processing of human mammary tissue and determination of total cell yield

Mammary gland tissue from healthy women undergoing reduction mammoplasty at the Nymphenburg Clinic for Plastic and Aesthetic Surgery was obtained in accordance with the regulations of the ethics committee of the Ludwig-Maximilians-University, Munich, Germany (proposal 397-12). Reduction mammoplasties used for this study are depicted in supplementary table S3 together with age and parity of the donor. Fresh breast tissue was transported on ice and processed immediately. Working on one small tissue piece at a time, the tissue was minced into 2-3 mm^3^ pieces using scalpels. Subsequently, minced tissue corresponding to roughly 5 mL was distributed to 50 mL falcons containing 20 mL tissue digestion buffer (DMEM/F12 w/o phenol red, 2% w/v BSA, 10 mM HEPES, 2 mM glutamine, 100 U/ml penicillin-streptomycin) supplemented with 1 μg/mL insulin (Sigma, I6634), 100 U/mL hyaluronidase (Sigma, H3506) and either 300 U/mL collagenase (Sigma, C9407) (protocol A) or 800 U/mL collagenase (protocol B and C). Depending on the amount of collected breast tissue per reduction mammoplasty sample, the number of 50 mL flacons filled with minced breast tissue and tissue digestion buffer differed but was never less than three. Enzymatic dissociation was performed at 37°C with vertical shaking at 10 rpm (protocol C) or 100 rpm (protocol A and B). After 3 hours (protocol A) or 16 hours (protocol B and C) completeness of the tissue digest was assessed by microscopic examination. The falcons were then filled up with an additional 20 mL of washing buffer (DMEM/F12 w/o phenol red, 10mM HEPES, 2mM glutamine, 100 U/ml penicillin-streptomycin) and cells were pelleted by centrifugation at 300 x g for 5 min. The fatty supernatant and overlaying medium were removed by aspiration and the pellets, containing epithelial fragments and stromal single cells, were washed once again with 25 mL washing buffer. For generation of a single-cell suspension, the epithelial fragments within the pellet were further dissociated using first 0,15% Trypsin-EDTA (Thermo Fisher, 25300054 mixed with 25200056) for 5 min followed by a 3 min treatment with 5 mg/mL dispase (Stem Cell Technologies, 07913) and 1 mg/mL DNase I (Sigma, 11284932001) and subsequently filtered through a 40 μm strainer. The total amount of cells isolated with dissociation protocols A, B and C was determined by manual counting of a defined single cell suspension volume and subsequent extrapolation for the total volume of dissociation suspension. Manual counting was performed using a Neubauer Improved chamber and the counting procedure was repeated three times for each sample. The pellets of single cells or epithelial fragments were resuspended in freezing medium (washing buffer, 10% DMSO, 50% FCS, 100 U/ml penicillin-streptomycin) at a cell density of roughly 5-7 x 10^6^ cells/mL and transferred to −80°C before being moved to liquid nitrogen for long term storage.

### Assessment of fragment morphology and measurement of fragment size

Brightfield images of epithelial fragments isolated with enzymatic dissociation protocols A, B and C from reduction mammoplasty samples M38, M47, M48, M49 and M50 were taken in 5x or 10x magnification. Fragment size was determined by measuring the area using the acquired brightfield images and ImageJ software. In total 1228 fragments (425 from protocol A, 400 from protocol B, 403 from protocol C) of reduction mammoplasties M48, M49 and M50 were analysed.

### Flow cytometry analysis and calculation of percentages and total numbers of epithelial and stromal fractions

Single-cell suspensions were generated from frozen fragments as described above and stained in a 0,1% BSA solution with CD31-PB, CD45-V450, CD49f-PE, EpCAM-FITC and CD10-APC. Prior to analysis, 7AAD (BD Bioscience, 559925) was added to distinguish dead and live cells. Cells were analysed on a FACSAriaTM III cell sorter (BD Biosciences) using a 100 μm nozzle and the FACSDivaTM 6.0 Software. For further analysis FLOWJo_V10 Software (FlowJo LLC) was used. A list of antibodies and dilutions used for flow cytometry is provided in supplementary table 4. For the determination of the percentages of different stromal and epithelial fractions isolated with enzymatic dissociation protocols A, B and C from reduction mammoplasties M46, M47, M48, M49, and M50, 1×10^6^ events were recorded for each sample and the number of viable, single cells was determined based on the gating strategy shown in Fig.S2A. Setting this number as 100 percent, the percentages of the three stromal (stroma 1, 2 and 3) and two epithelial populations (basal/myoepithelial and luminal) was determined based on the gating strategy show in Fig.2A. A list of percentage numbers of isolated stromal and epithelial populations is provided in supplementary file 1. The percentages of viable stromal and epithelial fractions were subsequently applied onto the total amount of isolated cells for each dissociation protocol and reduction mammoplasty sample to determine the total cell count per population and falcon of dissociation suspension.

### Immunofluorescence of epithelial fragments

A suspension containing epithelial fragments from reduction mammoplasty samples M48 and M49 was mixed with acidified rat tail Collagen I (Corning, 356236) and neutralizing solution (11x PBS, 550mM HEPES, comprising 1/10th of the volume of collagen) to a final concentration of 1,3 mg/mL collagen and the mixture was transferred into a 24-well plate as previously described ^21^. After polymerization of the collagen, the gels were washed once with PBS and then fixed with 4% paraformaldehyde (VWR, 43368) for 10 min. Cells were permeabilized with 0,2 % Triton X-100 (Sigma, T8787) and blocked with 10% donkey serum (GenTex, GTX73205) in 0.1% BSA. A list of primary and secondary antibodies used for staining is given in supplementary table S5 and S6. DAPI (Sigma, D9542) was used for visualization of cell nuclei. In total 23 fragments were imaged and representative images are shown in Fig.1D.

### Sample preparation for single cell RNA sequencing

Samples frozen as a mixture cellular fragments and single cells were used for single cell RNA sequencing experiments. After thawing, a single-cell suspension was generated as described above using trypsin and dispase treatment. Dead cells were eliminated via MACS using the Dead Cell Removal Kit from Milteny Biotec (130-090-101). Subsequently, cells were resuspended in 0.05% UltraPure BSA (Thermo Fisher Scientific, AM2616) and counted manually using a Neubauer Improved chamber. Cell concentration was adjusted to 800 cells/μL and equal numbers of cells were subjected to library preparation, which was performed according to the 10X Chromium Single Cell 3’ kit v3 instructions.

### Library preparation, sequencing and data processing

Library preparation was performed with the 10X Chromium single-cell kit using version 3 chemistry according to the instructions in the kit. The libraries were then pooled and sequenced on a NovaSeq600 S2 flow cell. Read processing was performed using the 10X Genomics workflow using the Cell Ranger Single-Cell Suite version 3.0.2. Samples were demultiplexed using barcode assignment and UMI quantification. The reads were aligned to the hg19 reference genome using the pre-built annotation package obtained from the 10X Genomics website (https://support.10xgenomics.com/single-cell-gene-expression/software/pipelines/latest/advanced/references). All lanes per sample were processed using the ‘cell ranger count’ function. The output from different lanes was then aggregated using ‘cellranger aggr’ with -normalize set to ‘none’.

### Quality control and data pre-processing

All downstream analysis was conducted using R version 4.0.0. Barcodes identified as containing low counts of unique molecular identifiers (UMIs) likely resulting from ambient RNA were removed using the function “emptyDroplets” from the *DropletUtils* package. Barcodes arising from single droplets containing cells were then filtered to ensure that cleaned barcodes contained at least 1000 UMIs and that the percentage of mitochondrial genes compared to overall annotated genes were not higher than 3x the median absolute deviation. Overall, 3,586 cells were obtained for Protocol A, 2,809 cells were obtained from Protocol B and 4,796 cells were obtained for Protocol C. Following filtering, cleaned barcodes were normalized and logged using the “computeSumFactors” from *scran* version 1.16.0 and “logNormCounts” from the *scater* package version 1.16.1. Principal component analysis was then performed on the normalized cells using *PCA tools* version 2.0.0 followed by uniform manifold approximation and projection graphing (*umap* version 0.2.6.0) and clustering based on generated igraphs and Louvain clustering conducted using *scran*. Overall, 14 clusters were identified. One cluster (C13), however, contained cells with low UMI counts and did not uniformly express markers associated with any mammary subpopulations and was therefore excluded (Fig. S7). For ease of interpretability and graphical representation of clusters arising from different cell subpopulations, we down-sampled our cell numbers to ensure 1,700 cells per digest type. PCA, UMAP and clustering analysis revealed 12 clusters for the equal numbers of subset cells per digest, which were then utilized for downstream analysis. Further sub-clustering was performed for certain cell types such as those annotated as immune cells. Plots of interesting genes were generated using either *ggplot2* or *pheatmap* packages with custom colours generated by the *RcolourBrewer* package.

### Differential gene expression analysis and GO term annotation

Differentially Expressed Genes (DEGs) were identified between subsetted groups by firstly generating a pseudo bulk signature for each cell type within the two dissociation duration groups of 3-hour dissociation and 16-hour dissociation using the “aggregateAcrossCells” function from the *scater* package. For the overall dissociation comparisons cells within each dissociation duration group, averaged cell type counts were converted to a DGEList, filtered for lowly expressed genes and normalized using functions within the package *edgeR* version 3.30.3. The average counts for the 16-hour dissociated cells were then taken away from the from the 3-hour dissociated cell counts and genes with a fold change of 1.5 were noted as being differentially expressed overall. DEGs were visualised as MA plots using the “ggplot” function from the ggplot2 package. DEGs identified from all cells in the 3-hour dissociation in comparison to 16-hour dissociation with a fold change greater than 1.5 had gene set enrichment analysis performed on them for gene ontology (GO) terms associated with biological process and molecular functions using the *topGO* package version 2.40.0. Similarly, cell type specific DEGs were identified by removing lowly expressed genes and normalizing between the average counts for the same cell type and determining genes that had a fold change greater than 2.0 between the fast and slow digests, which were then interrogated for molecular function GO terms. A list of differentially expressed genes and GO term annotations are provided in supplementary files 2-15.

### Determining enrichment for gene expression signatures across individual cells

Overall expression of gene signatures identified from highly expressed DEGs in all cells was examined using the “AddModuleScore” function from the *Seurat* package version 3.1.5. Five randomly selected genes were used as a control for these genes to determine a “oxidative stress” signature score for each of the cells. The resulting score is unitless but is indicative of signature enrichment when compared between cell type groups. A list of genes used for the “oxidative stress” gene set is provided in Table S2.

## Data availability

The authors declare that all data supporting the findings of this study are available within the article, its supplementary figures, or from the corresponding author upon reasonable request. The RNA sequencing data will be deposited in the Gene Expression Omnibus (GEO) database and released upon publication. All computational analyses were performed in R (Version 4.0.0) using standard functions unless otherwise indicated. All Codes used will be available online at https://github.com/.

## Acknowledgements

We thank the Helmholtz sequencing core and particularly Sandy Loesecke for conducting the sequencing. We thank Thomas Walzthoeni for bioinformatics support provided at the Bioinformatics Core Facility, ICB, Helmholtz Center Munich. Our thanks are extended to Karsten Bach (Department of Pharmacology, University of Cambridge) and Malte Lücken (Institute of Computational Biology, ICB, Helmholtz Center Munich) for direction in scRNA-analysis. We thank Anika Böttcher (Institute of Diabetes and Regeneration Research, IDR, Helmholtz Center Munich) for sharing experimental expertise. We are grateful to Magdalena Götz (Institute for stem cell research, ISF, Helmhotz Center Munich) and Stefan Stricker (Institute for stem cell research, ISF, Helmholtz Center Munich) for helpful discussions. We thank all members of the Scheel group and the Khaled group for support, constructive discussions and for sharing experimental expertise. Many thanks to all the participants of this study. This work was funded by a Max Eder Grant of the German Cancer Aid Foundation (Deutsche Krebshilfe 110225 to C.H.S.). A.J.T. was funded by the Helmholtz postdoctoral fellowship program and a BBSRC project Grant (BB/S006745/1) to W.T.K.

## Author Contributions

L.K.E. conceived the project and designed experiments. L.K.E., A.J.T., H.M.G. and C.G. processed reduction mammoplasties. L.K.E. performed flow cytometry experiments and immunofluorescence experiments and together with M.S. and I.K. prepared the library for scRNA-seq. A.J.T. conducted bioinformatic analysis. Data were interpreted by L.K.E., A.J.T., H.M.G., A.R.B., W.T.K. and C.H.S. The manuscript was prepared by L.K.E., A.J.T. and C.H.S. and critically reviewed by all authors.

## Figure Legends

**Figure S1:**
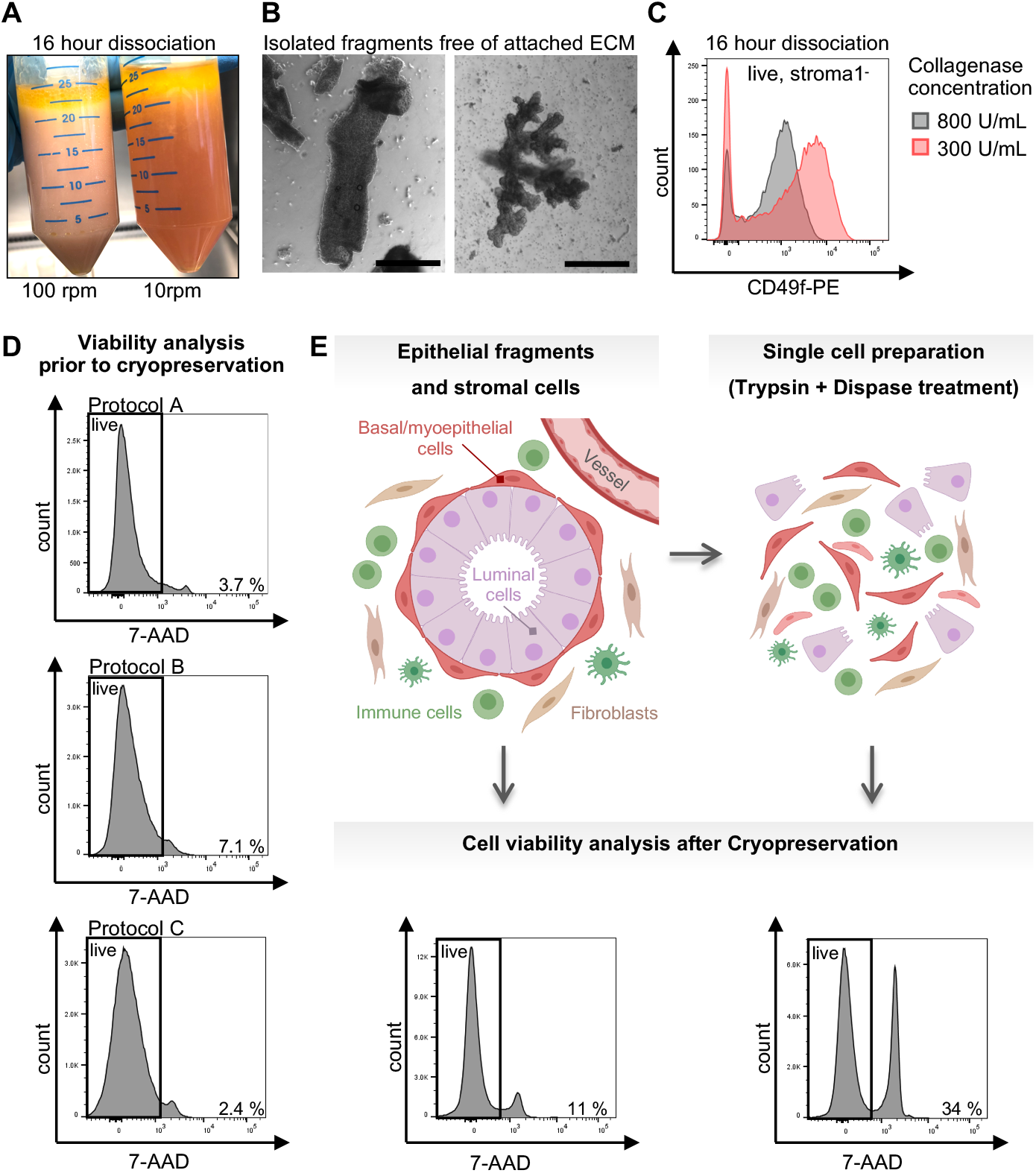
Freezing strategy of primary mammary cells influences cells viability. Enzymatic dissociation was considered successful if **(A)** no remaining tissue clumps were visible in the dissociation suspension, which was clearly separated from the overlaying fat layer (Reduction mammoplasty: M48) and **(B)** isolated fragments were free of any attached extracellular matrix. Scale bar = 500 μm. (Reduction mammoplasty: M48). **(C)** CD49f was downregulated after a 16-hour enzymatic dissociation process at 10 rpm using 800 instead of 300 U/mL collagenase as determined via flow cytometry after exclusion of doublets, dead cells (live=7-AAD^−^) and the stroma1 population (stroma1^−^=CD31^−^/CD45^−^). Reduction mammoplasty: M46 **(D)** Representative flow cytometry-based viability analysis prior to cryopreservation using live-dead marker 7-AAD showed high viability after enzymatic dissociation protocols A, B and C. (7-AAD^−^ = live), Reduction mammoplasty: M48. **(E)** Mammary cells frozen as a mixture of epithelial fragments and single cells showed higher cell viability after thawing (left panel) than cells that had been subjected to trypsinization and dispase treatment and were frozen as a single cell suspension (right panel). Flow cytometry-based cell viability analysis was performed after exclusion of doublets using the live-dead marker 7-AAD (7-AAD^−^=live). Reduction mammoplasty: M44. Created with BioRender.com.

**Figure S2:**
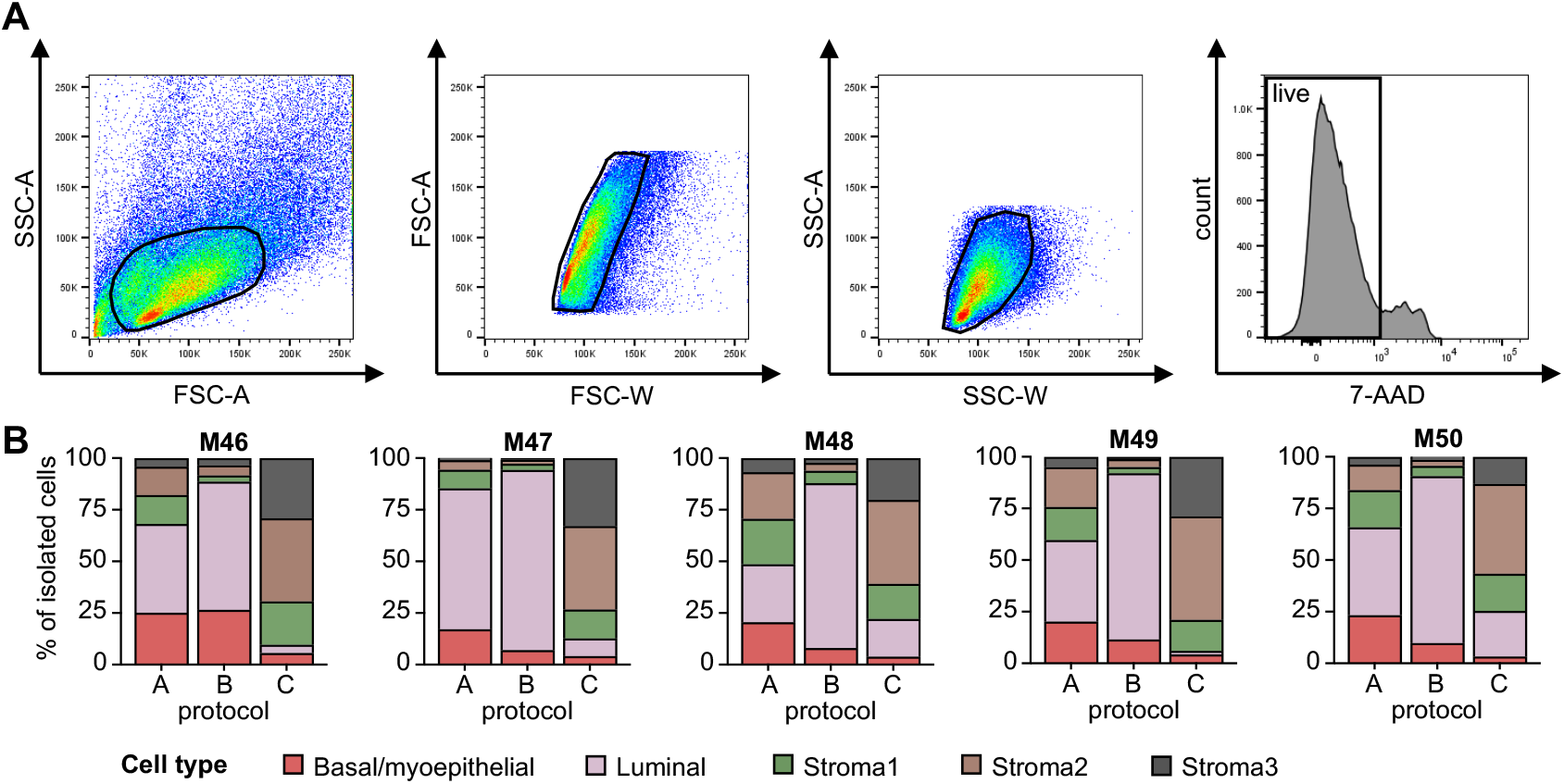
Proportions of stromal and epithelial populations isolated with different dissociation protocols. **(A)** Flow cytometry gating strategy for exclusion of doublets and dead cells (live=7-AAD^−^). **(B)** Proportions of stromal and epithelial populations isolated with enzymatic dissociation protocol A, B and C based on flow cytometry analysis as shown in Fig.2A Reduction mammoplasties: M46, M47, M48, M49, M50.

**Figure S3:**
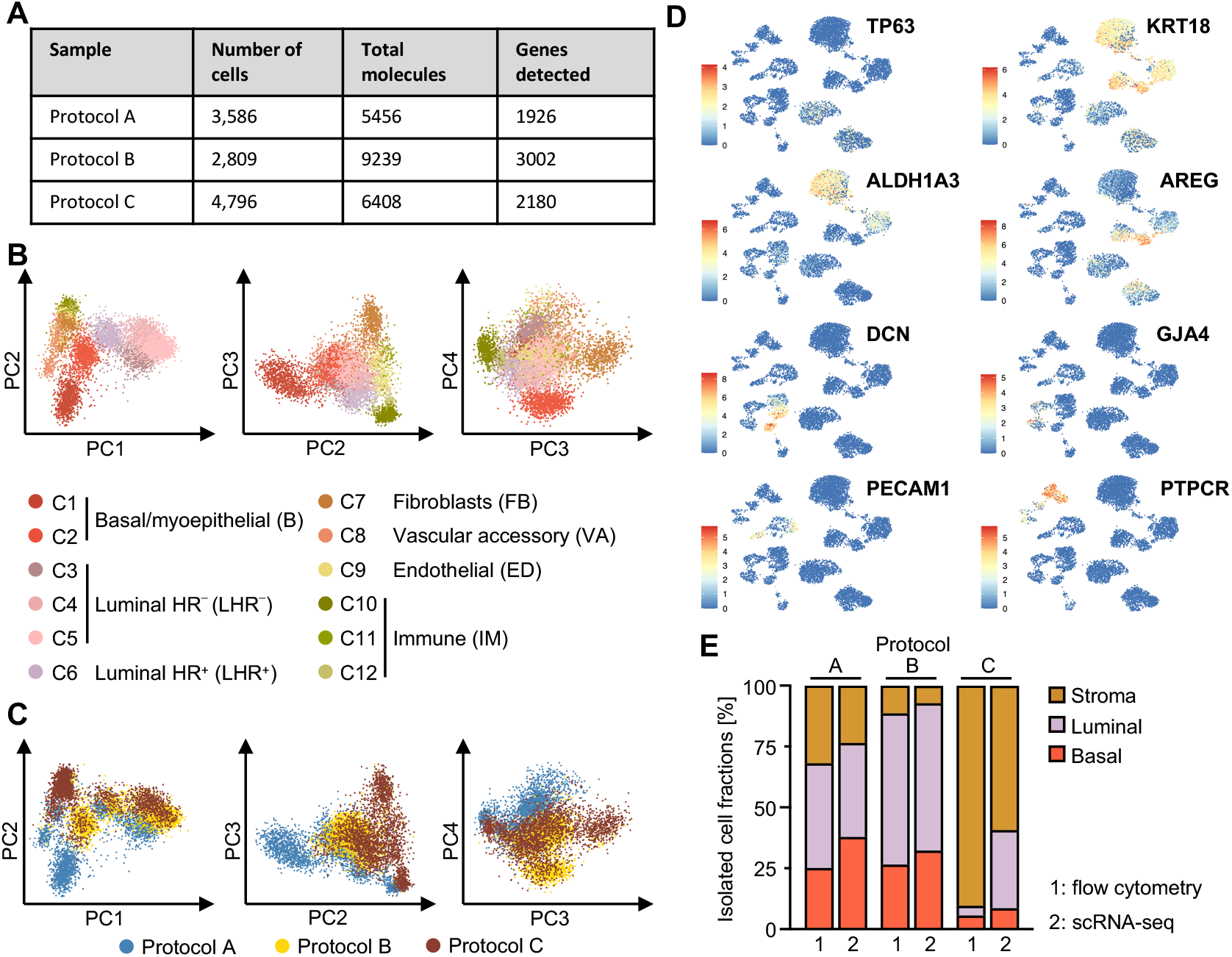
Single-cell RNA sequencing analysis of cells isolated with enzymatic dissociation protocols A, B, and C. **(A)** Table summarizing number of cells sequenced from dissociation protocol A, B and C of reduction mammoplasty M46 and quality control criteria per sample (number of unique molecules, genes detected and number of reads). **(B)** The same 9,000 cells shown in UMAP plots in Fig.3 A,B were displayed in principal component analysis plots coloured by clusters derived from UMAP plot (Fig.1A) and **(C)** by dissociation protocol. **(D)** UMAP plots coloured by normalized log-transformed expression of cell type specific marker genes. **(E)** Proportion of epithelial and stromal cell types isolated with enzymatic dissociation protocols A, B, and C of reduction mammoplasty M46 and detected either via flow cytometry (1) or scRNA-seq. (2).

**Figure S4:**
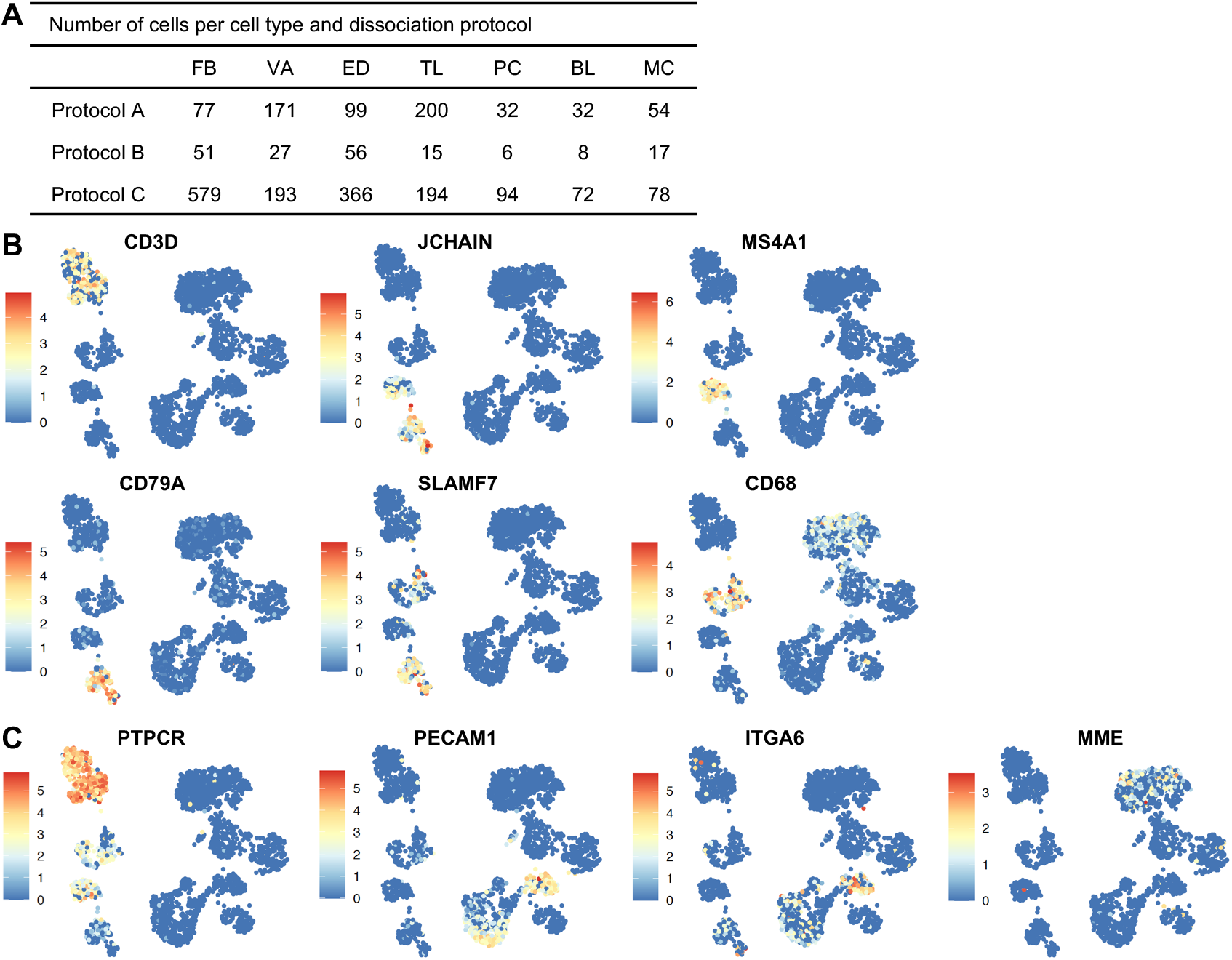
Marker gene analysis to identify putative identities of mammary stromal cell clusters. **(A)** Table summarizing number of cells making up the stromal subclusters derived from dissociation protocol A, B, and C. **(B)** UMAP plots showing the stromal subclusters (SC1-SC12) coloured by normalized log-transformed expression of selected marker genes used for identification of different immune cell types. **(C)** UMAP plot showing the stromal subclusters (SC1-SC12) coloured by normalized log-transformed expression of *PTPCR*, *PECAM1*, *ITGA6* and *MME*.

**Figure S5:**
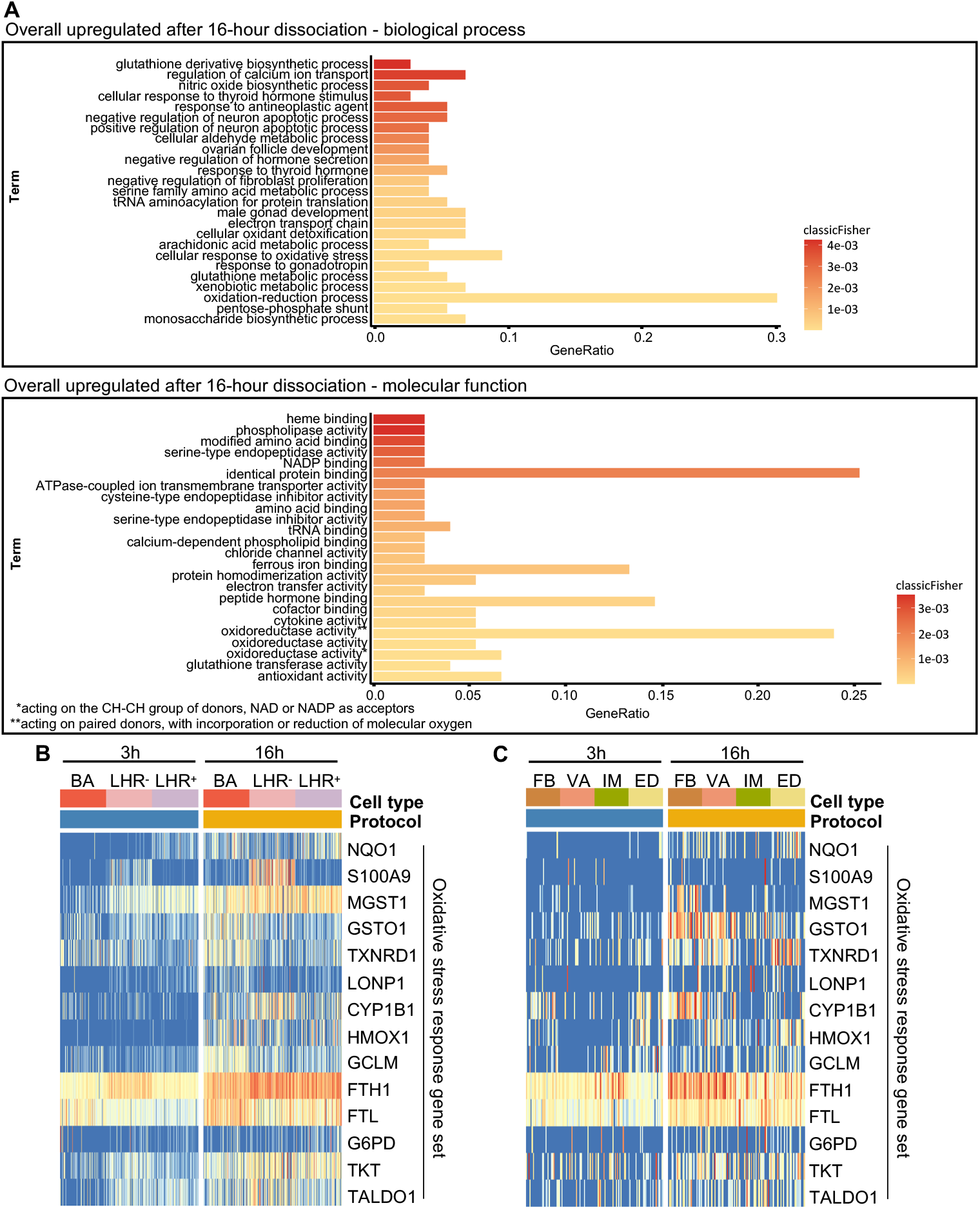
GO term analysis of genes upregulated after a 16-hour compared to a 3-hour dissociation reveals an oxidative stress response. **(A)** GO term analysis of 60 genes upregulated (log fold change > 1.5) in cells after a 16-hour (protocol B and C) compared to a 3-hour dissociation (protocol A). List of GO term biological process (upper graph) and molecular function (lower graph) is displayed as a bar chart in which the colour represents the p-value of the term and the bar length is proportional to the overlapping genes. **(B, C)** Heatmaps showing genes selected for oxidative stress response gene set in epithelial populations **(B)** and stromal populations **(C)**. Upper bars represent cell type/cluster and dissociation protocol origin of cells, 3-hour (protocol A) and 16-hour dissociation (protocol B and C). 150 randomly selected cells of epithelial populations and 30 of stromal populations are displayed for better visualization. Colour scale represents log-transformed and normalized counts scaled to a maximum of 1 per row.

**Figure S6:**
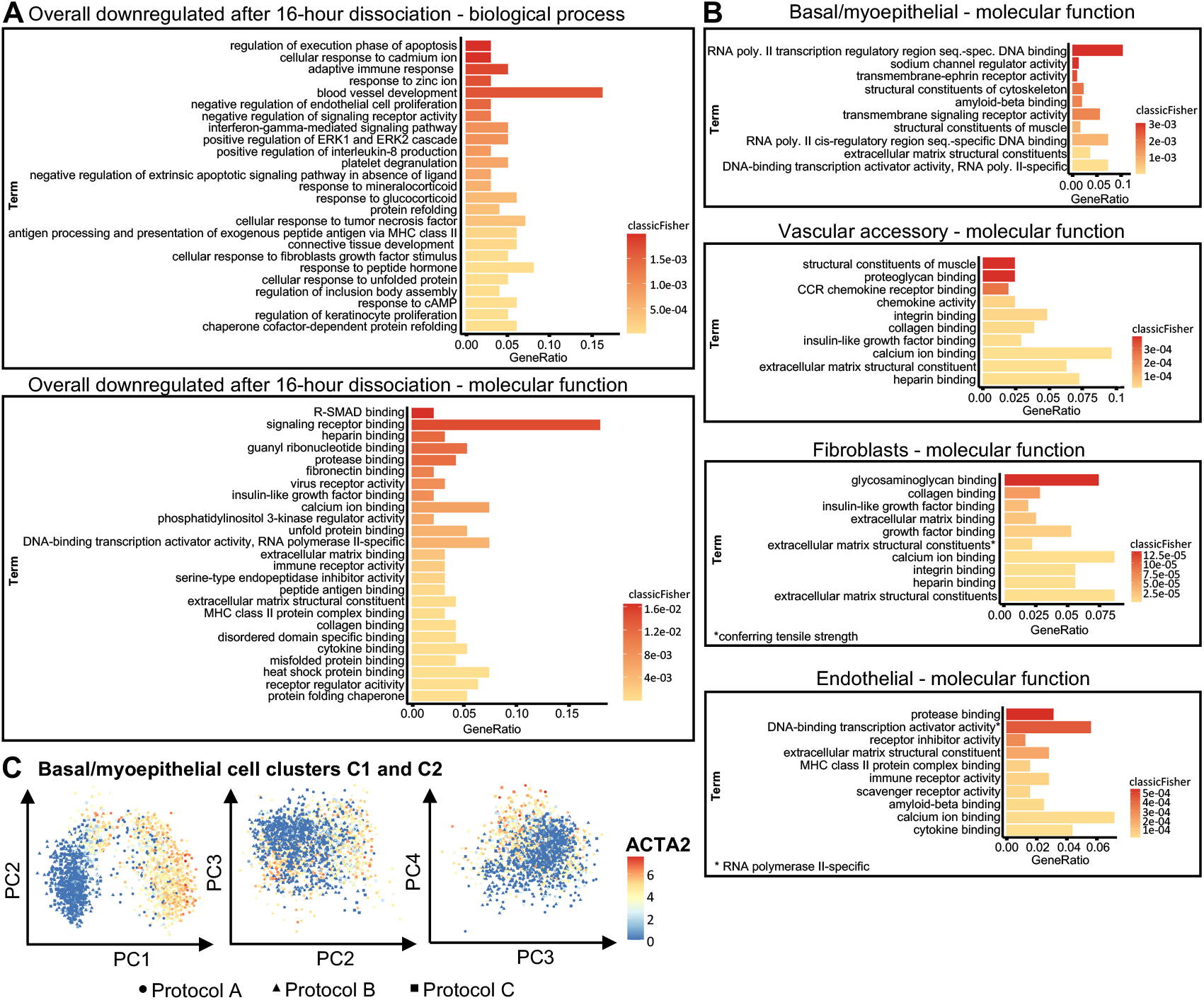
GO term analysis of genes downregulated after a 16-hour compared to a 3-hour dissociation reveals downregulation of lineage specific marker genes. **(A)** GO term analysis of 132 genes downregulated (log fold change>1.5) in cells after a 16-hour (protocol B and C) compared to a 3-hour dissociation (protocol A). Selected significantly enriched GO terms (P<0.01) are displayed. List of GO term biological process (upper graph) and molecular function (lower graph) is displayed as a bar chart in which the colour represents the p-value of the term and the bar length is proportional to the overlapping genes. **(B)** GO term analysis of genes downregulated (log fold change>2.0) after a 16-hour (protocol B and C) compared to a 3-hour dissociation (protocol A) within the basal/myoepithelial, vascular accessory, fibroblast and endothelial population. For each population, a list of GO term molecular functions is displayed as a bar chart in which the colour represents the p-value of the term and the bar length is proportional to the overlapping genes. **(C)** Basal/myoepithelial cells derived from the 16-hour (protocol B and C, cluster C2) and from the 3-hour dissociation (protocol A, cluster C1) are displayed by principal component analysis coloured by normalized log-transformed expression of marker gene *ACTA2*.

**Figure S7:**
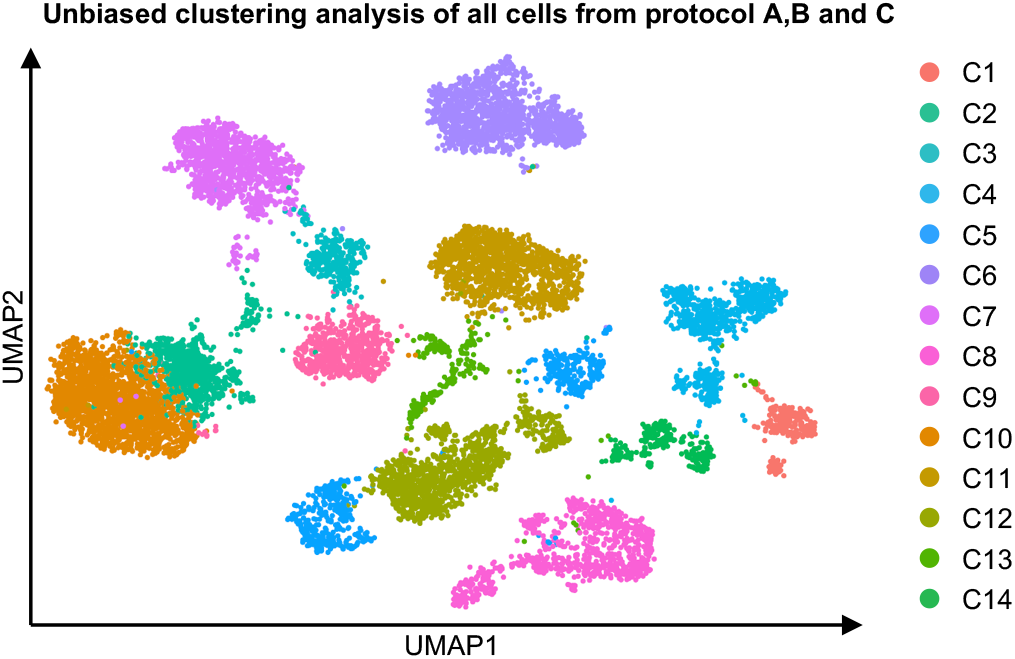
Clustering analysis of all cells isolated with different dissociation protocols. All cells from dissociation protocol A, B and C of reduction mammoplasty M46 were visualized as 14 clusters in a UMAP plot and coloured by cluster. Cluster 13 (C13) contained cells with low UMI counts and did not uniformly express markers associated with any mammary subpopulations and was therefore excluded.

## Supplementary Tables

**Table S1:** Enzymatic dissociation of human breast tissue.

| Duration (hours) | Agitation speed | Enzyme concentrations | Reference |
| --- | --- | --- | --- |
| 16-18 | n.a. | 300 U/mL Collagenase, 100U/mL Hyaluronidase | Emerman, J. T. & Wilkinson, D. A. **Routine culturing of normal, dysplastic and malignant human mammary epithelial cells from small tissue samples.** *Vitr. Cell. Dev. Biol.* (1990)<sup>63</sup> |
| 24 | gentle | 450 U/mL Collagenase | Ögmundsdóttir, H. M. et al. **Effects of lymphocytes and fibroblasts on the growth of human mammary carcinoma cells studied in short-term primary cultures.** *Vitr. Cell. Dev. Biol.-Anim.* (1993)<sup>64</sup> |
| 16-18 | gentle | 300 U/mL Collagenase, 100U/mL Hyaluronidase | Stingl, J., Eaves, C. J., Kuusk, U. & Emerman, J. T. **Phenotypic and functional characterization in vitro of a multipotent epithelial cell present in the normal adult human breast.** *Differentiation* (1998)<sup>65</sup> |
| 24 | n.a. | 900 U/mL Collagenase | Rønnov-Jessen, L., Villadsen, R., Edwards, J. C. & Petersen, O. W. **Differential expression of a chloride intracellular channel gene, CLIC4, in transforming growth factor-β1-mediated conversion of fibroblasts to myofibroblasts.** *Am. J. Pathol.* (2002)<sup>66</sup> |
| 5-8 | n.a. | 150 U/mL Collagenase, 50U/mL Hyaluronidase | Lim, E. et al. **Aberrant luminal progenitors as the candidate target population for basal tumor development in BRCA1 mutation carriers.** *Nat. Med.* (2009)<sup>55</sup> |
| overnight | 8 rpm | 200 U/mL Collagenase, 100U/mL Hyaluronidase | Labarge, M. A., Garbe, J. C. & Stampfer, M. R. **Processing of human reduction mammoplasty and mastectomy tissues for cell culture.** *J. Vis. Exp.* 1–7 (2013)<sup>19</sup> |
| 6-8 | 300-500 rpm | 2mg/mL Collagenase | Hines, W. C., Su, Y., Kuhn, I., Polyak, K. & Bissell, M. J. **Sorting out the FACS: A devil in the details.** *Cell Rep.* 6, 779–781 (2014)<sup>13</sup> |
| 18-24 | 80 rpm | 1mg/mL Collagenase |  |
| 6-8 | 300 rpm | 250U/mL Collagenase, 1400U/mL Hyaluronidase | Zubeldia-Plazaola, A. et al. **Comparison of methods for the isolation of human breast epithelial and myoepithelial cells.** *Front. Cell Dev. Biol.* 3, 1–9 (2015)<sup>18</sup> |
| overnight | 50 rpm | 200U/mL collagenase, 100U/mL hyaluronidase |  |
| overnight | n.a. | 2mg/mL collagenase | Nguyen, Q. H. et al. **Profiling human breast epithelial cells using single cell RNA** **sequencing identifies cell diversity.** *Nat. Commun.* (2018)<sup>6</sup> |
| 2 | n.a. | 1mg/mL collagenase | Rosenbluth, J. M. et al. **Organoid cultures from normal and cancer-prone human breast tissues preserve complex epithelial lineages.** *Nat. Commun.* (2020)<sup>67</sup> |

Summary of enzymatic dissociation protocols for human breast tissue samples from selected publications. The dissociation parameters duration, agitation speed and enzyme concentrations are displayed. n.a.: data not available.

**Table S2:** Human Reduction mammoplasties.

| Reduction mammoplasty donor | Age (years) | Parity |
| --- | --- | --- |
| M38 | 47 | 2 |
| M40 | 65 | 1 |
| M43 | 54 | 2 |
| M44 | 19 | 0 |
| M45 | 54 | 1 |
| M46 | 51 | 1 |
| M47 | 55 | 2 |
| M48 | 49 | 1 |
| M49 | 68 | 1 |
| M50 | 58 | 1 |

**Table S3:**
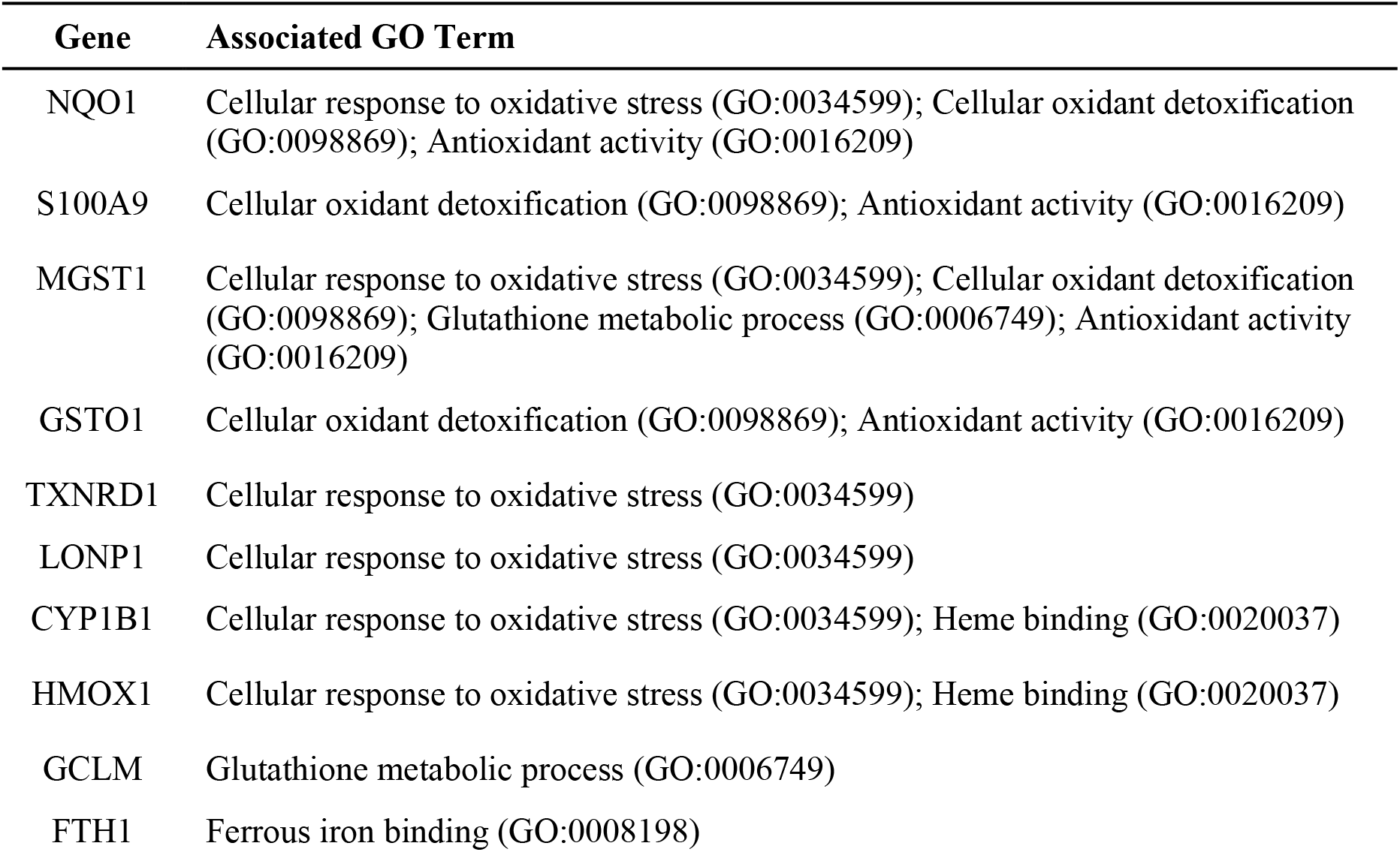
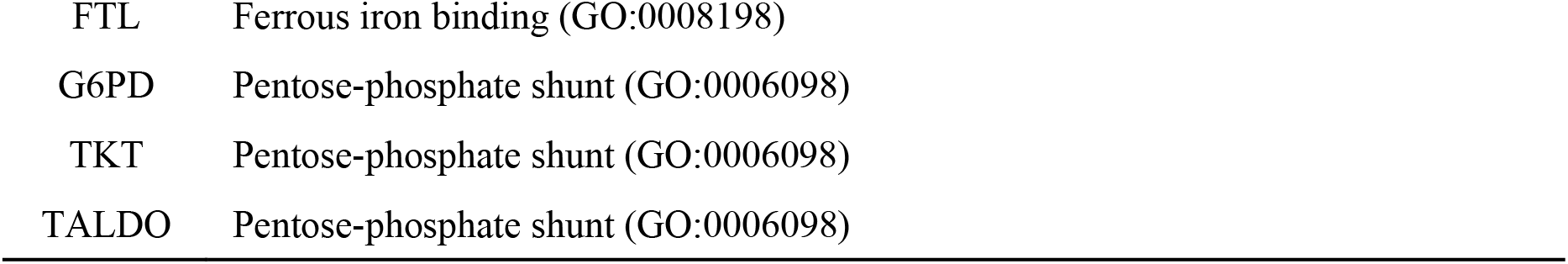
Stress response gene set.

**Table S4:** Antibodies used for flow cytometry.

| Epitope [Clone] | Conjugation | Host | Volume (μL) * | Catalog number | Supplier |
| --- | --- | --- | --- | --- | --- |
| 7-AAD | - | - | 2 | 559925 | BD, Heidelberg, Germany |
| CD10 [HI10A] | APC | Mouse | 2.5 | 312210 | Biozol, Eching, Germany |
| CD31 [WM59] | PB | Mouse | 0.5 | 303114 | Biozol, Eching, Germany |
| CD326/EPCAM [VU-1D9] | FITC | Mouse | 5 | GTX798 49 | Biozol, Eching, Germany |
| CD45 [HI30] | V450 | Mouse | 0.5 | 560367 | Biozol, Eching, Germany |
| CD49f [GOH3] | PE | Rat | 2.5 | 555736 | BD, Heidelberg, Germany |

\* Used to stain $1 \times 10^6$ cells

**Table S5:** Primary antibodies.

| Epitope [Clone] | Host | Dilution | Catalog number | Supplier |
| --- | --- | --- | --- | --- |
| Alpha smooth muscle actin | Rabbit | 1/100 | ab5964 | Abcam, Cambridge, UK |
| Cytokeratin 8/18 [5D3] | Mouse | 1/250 | DLN-10750 | Dianova, Hamburg, Germany |
| ZO-1 | Mouse | 1/100 | 339100 | Life Technologies, Darmstadt, Germany |

**Table S6:** Secondary antibodies.

| Host/Isotype | Species reactivity | Conjugation | Dilution | Catalog number | Supplier |
| --- | --- | --- | --- | --- | --- |
| Donkey/IgG | Mouse | Alexa 488 | 1/250 | A-21202 | Life Technologies, Darmstadt, Germany |
| Donkey/IgG | Rabbit | Alexa 546 | 1/250 | A-10040 | Life Technologies, Darmstadt, Germany |

## Description of additional Supplementary Files

**File name: Supplementary File 1**

Description: Percentage numbers of mammary cell populations identified via flow cytometry

**File name: Supplementary File 2**

Description: Differentially expressed genes between cells derived from protocol A (3-hour dissociation) and from protocol B/C (16-hour dissociation)

**File name: Supplementary File 3**

Description: Differentially expressed genes between basal/myoepithelial (BA) cells derived from protocol A (3-hour dissociation) and from protocol B/C (16-hour dissociation)

**File name: Supplementary File 4**

Description: Differentially expressed genes between Luminal HR- (LHR-) cells derived from protocol A (3-hour dissociation) and from protocol B/C (16-hour dissociation)

**File name: Supplementary File 5**

Description: Differentially expressed genes between Luminal HR+ (LHR+) cells derived from protocol A (3-hour dissociation) and from protocol B/C (16-hour dissociation)

**File name: Supplementary File 6**

Description: Differentially expressed genes between fibroblasts (FB) derived from protocol A (3-hour dissociation) and from protocol B/C (16-hour dissociation)

**File name: Supplementary File 7**

Description: Differentially expressed genes between vascular accessory (VA) cells derived from protocol A (3-hour dissociation) and from protocol B/C (16-hour dissociation)

**File name: Supplementary File 8**

Description: Differentially expressed genes between endothelial (ED) cells derived from protocol A (3-hour dissociation) and from protocol B/C (16-hour dissociation)

**File name: Supplementary File 9**

Description: Differentially expressed genes between immune (IM) cells derived from protocol A (3-hour dissociation) and from protocol B/C (16-hour dissociation)

**File name: Supplementary File 10**

Description: GO terms for genes upregulated after a 16-hour dissociation (protocol B/C) compared to a 3-hour dissociation (protocol A)

**File name: Supplementary File 11**

Description: GO terms for genes downregulated after a 16-hour dissociation (protocol B/C) compared to a 3-hour dissociation (protocol A)

**File name: Supplementary File 12**

Description: GO terms for differentially expressed genes between basal/myoepithelial (BA) cells derived from protocol A (3-hour dissociation) and from protocol B/C (16-hour dissociation)

**File name: Supplementary File 13**

Description: GO terms for differentially expressed genes between vascular accessory (VA) cells derived from protocol A (3-hour dissociation) and from protocol B/C (16-hour dissociation)

**File name: Supplementary File 14**

Description: GO terms for differentially expressed genes between fibroblasts (FB) derived from protocol A (3-hour dissociation) and from protocol B/C (16-hour dissociation)

**File name: Supplementary File 15**

Description: GO terms for differentially expressed genes between endothelial (ED) cells derived from protocol A (3-hour dissociation) and from protocol B/C (16-hour dissociation)

